# Postsynaptic plasticity of cholinergic synapses underlies the induction and expression of appetitive memories in *Drosophila*

**DOI:** 10.1101/2021.07.01.450776

**Authors:** Carlotta Pribbenow, Yi-chun Chen, Michael-Marcel Heim, Desiree Laber, Silas Reubold, Eric Reynolds, Isabella Balles, Raquel Suárez Grimalt, Carolin Rauch, Jörg Rösner, Gregor Lichtner, Sridhar R. Jagannathan, Tania Fernández-d.V. Alquicira, David Owald

**Affiliations:** Institute of Neurophysiology, Charité – Universitätsmedizin Berlin, corporate member of Freie Universität Berlin and Humboldt-Universität zu Berlin, and Berlin Institute of Health, Charitéplatz 1, 10117 Berlin, Germany; Einstein Center for Neurosciences Berlin, Charitéplatz 1, 10117 Berlin; NeuroCure, Charité – Universitätsmedizin Berlin, corporate member of Freie Universität Berlin and Humboldt-Universität zu Berlin, and Berlin Institute of Health, Charitéplatz 1, 10117 Berlin, Germany; NWFZ, Charité – Universitätsmedizin Berlin, corporate member of Freie Universität Berlin and Humboldt-Universität zu Berlin, and Berlin Institute of Health, Charitéplatz 1, 10117 Berlin, Germany; Universitätsmedizin Greifswald, Department of Anesthesia, Critical Care, Emergency and Pain Medicine, Ferdinand-Sauerbruch-Straße, 17475 Greifswald, Germany

**Keywords:** learning and memory, postsynaptic plasticity, cholinergic synapses, *Drosophila*, mushroom bodies

## Abstract

In vertebrates, several forms of memory-relevant synaptic plasticity involve postsynaptic rearrangements of glutamate receptors. In contrast, previous work indicates that *Drosophila* and other invertebrates store memories using presynaptic plasticity of cholinergic synapses. Here, we provide evidence for postsynaptic plasticity at cholinergic output synapses from the *Drosophila* mushroom bodies (MBs). We find that the nicotinic acetylcholine receptor (nAChR) subunit α5 is required within specific MB output neurons (MBONs) for appetitive memory induction, but is dispensable for aversive memories. In addition, nAChR α2 subunits mediate memory expression and likely functions downstream of α5 and the postsynaptic scaffold protein Dlg. We show that postsynaptic plasticity traces can be induced independently of the presynapse, and that *in vivo* dynamics of α2 nAChR subunits are changed both in the context of associative and non-associative memory formation, underlying different plasticity rules. Therefore, regardless of neurotransmitter identity, key principles of postsynaptic plasticity support memory storage across phyla.

## Introduction

Changing the strength of defined chemical synaptic connections within associative networks is widely believed to be the basis of memory storage^1–3^. However, it is unclear and frequently debated to what degree underlying neurophysiological and molecular mechanisms are evolutionarily conserved. One main difference between vertebrates and invertebrates is that memory-storing synapses in vertebrates use glutamate as their primary transmitter, while those in invertebrates (at least for *Drosophila melanogaster* and *Sepia officinalis*) use acetylcholine^4–6^. Furthermore, it is widely established that invertebrate nervous systems utilize presynaptic plasticity for storing memories, while the degree to which postsynaptic plasticity can be used is largely unclear. In contrast, it is well established that storing information in vertebrates can depend on both pre- and postsynaptic mechanisms, including postsynaptic rearrangements of neurotransmitter receptors.

Detailed knowledge of the anatomical wiring and functional signaling logic of the *Drosophila* learning and memory centers, the mushroom bodies (MBs)^5,7–21^, allows one to address whether, despite the use of different neurotransmitter systems, memory storage modes are functionally comparable or evolutionarily conserved. The weights of Kenyon cells (KCs) to MB output neuron (MBON) synapses are modulated by dopaminergic neurons (DANs), which anatomically divide the MBs into at least 15 functional compartments, where information is stored on appetitive and aversive associations, in addition to non-associative learning such as the relative familiarity of an odor^5,7–12,15,22,23^.

Studies so far have identified several traits pointing towards presynaptic storage mechanisms within the KCs during memory formation^24–27^. Indeed, studies that have blocked neurotransmitter release from KCs during learning^28–31^ have brought postsynaptic contributions to synaptic plasticity into question.

In vertebrates, typically, postsynaptic long-term changes^2,32,33^ are mediated via NMDA-sensitive glutamate receptors (NMDAR) that induce (‘induction’^3^) an expression phase (‘expression’^3^) through changed glutamatergic AMPA receptor (AMPAR) dynamics in dependence of postsynaptic scaffolds like PSD-95^34^. Invertebrate nAChRs in principle could take over similar functions to their glutamatergic counterparts in vertebrates, despite their differing molecular characteristics^35^. Indeed, nAChRs are pentamers that can be composed of homomeric assemblies of α subunits or heteromeric combinations of different α and β subunits. The composition of subunits determines the physiological properties of the nAChRs^35–38^, and synaptic weights could, in theory, be adjusted through the exchange of receptor subunits or entire complexes.

Here, we capitalize on the genetic accessibility to individual output neurons of the MBs to directly test whether postsynaptic receptors play a role in memory storage. Derived from combined neurophysiological, behavioral, light microscopic and molecular approaches, our data are supportive of a sequential role for nAChR subunits in appetitive memory storage at the level of MBONs. Using artificial training protocols, we demonstrate that postsynaptic calcium transients can change in response to concurrent activation of dopaminergic neurons and application of acetylcholine, circumventing KC output. Blocking KC output during appetitive, but not aversive, learning abolishes memory performance. Moreover, specific knock-down of the α5 nAChR subunit, but none of the other six α subunits, in the M4/6 MBONs (also known as MBON-γ5β’2a, MBON-β’2mp, MBON-β2β’2a and MBON-β’2mp bilateral) – an output junction involved in coding appetitive and aversive memories^7,11,17^ – impairs immediate appetitive memories. Knock-down of α2 or α5, however, interferes with 3-hour appetitive memories, as does knock-down of the scaffold Discs large (Dlg). We report differential distribution of α subunits throughout the MB and demonstrate that signal recovery of GFP-tagged subunits (as measured through fluorescence signal recovered after photobleaching) is changed through plasticity protocols. In addition, postsynaptically expressed non-associative familiarity learning also depends on α5 and α2 signaling as well as α2 dynamics. We propose a temporal receptor model and speculate that, in *Drosophila*, nAChR subunits α5 and α2 take roles similar to NMDA and AMPA receptors in vertebrates for memory induction and expression, indicating that the general principle for postsynaptic plasticity independent of the neurotransmitter system used, could be conserved throughout evolution.

## Results

### Neurotransmitter release from Kenyon cells is required for appetitive learning

One key argument in favor of exclusively presynaptic memory storage mechanisms in *Drosophila* is based on experiments suggesting that blocking KC or KC subset output selectively during learning leads to unaltered or mildly changed memory performance^28–31^. If the postsynapse need not see the neurotransmitter during training, it would likely be dispensable for memory induction. We revisited such experiments and blocked KC output during T-maze training, exposing the animals either to sugar-odor or shock-odor pairings (Fig. 1 a-b, Supplementary Fig. 1a-b).

**Figure 1:**
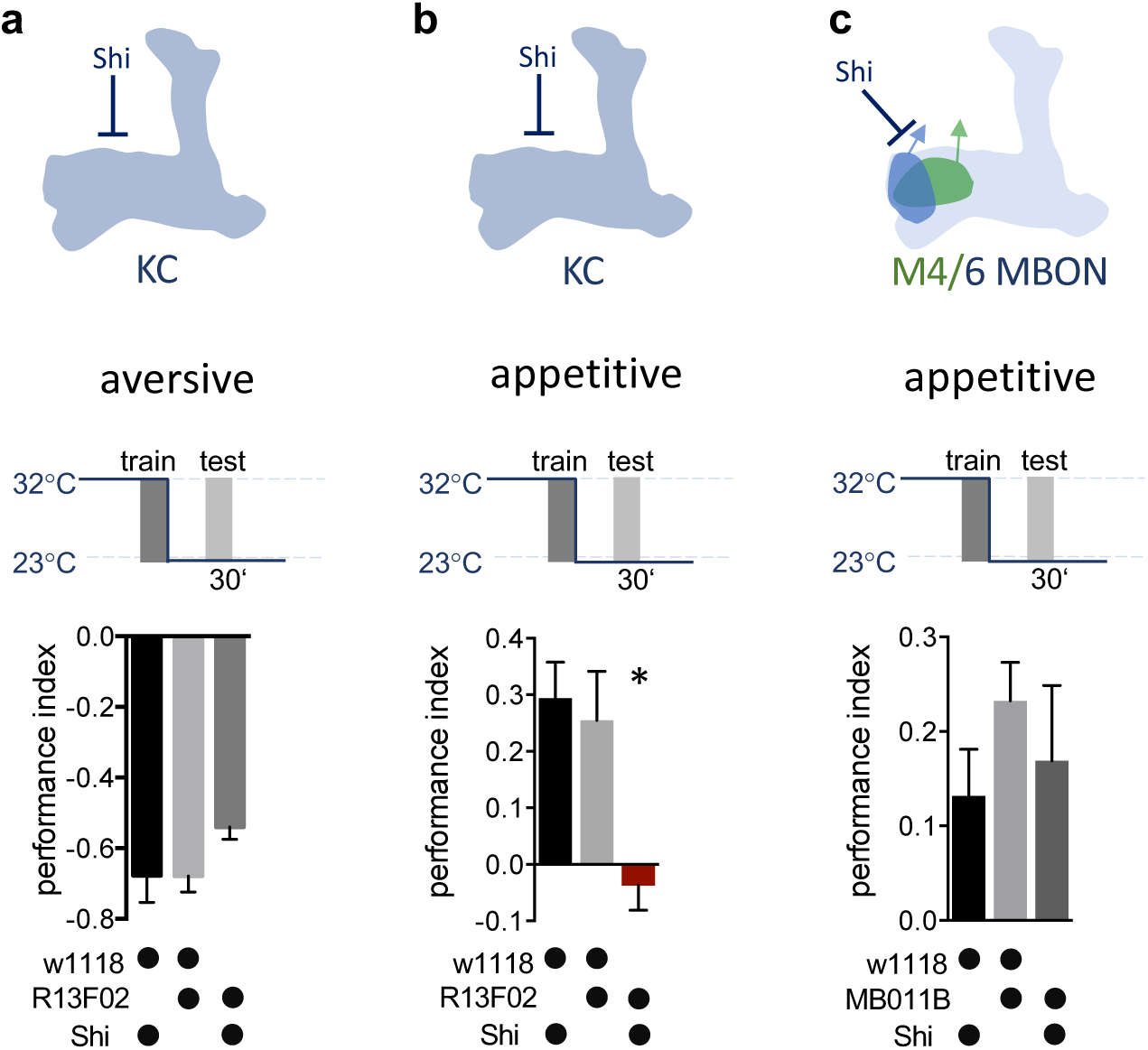
KC neurotransmitter release is required for the acquisition of appetitive memories. **a-c)** Flies expressing temperature-sensitive Shibire (Shi) within KCs or MBONs are trained at restrictive temperature (32°C), and subsequently placed at permissive temperature (23°C) throughout the consolidation and retrieval phase. Memory performance was tested 30 minutes after training at permissive temperature. Shi blocks neurotransmitter release at 32°C. **a)** Block of neurotransmitter release from KCs (driver line R13F02-Gal4) during training does not impact 30 min aversive memory performance. Bar graphs: mean ± SEM; n = 7 – 8; one-way ANOVA followed by Tukey’s test (p > 0.05). **b)** Block of neurotransmitter release from KCs (driver line R13F02-Gal4) during training impairs 30 min appetitive memory performance. Bar graphs: mean ± SEM; n = 10 – 16; Kruskal-Wallis followed by Dunn’s test (p < 0.05), * = p < 0.05. **c)** Block of neurotransmitter release from M4/6 MBONs (driver line MB011B) during training does not impact 30 min appetitive memory performance. Bar graphs: mean ± SEM; n = 14 – 24; one-way ANOVA followed by Tuckey’s test (p > 0.05). Also see Supplementary Fig. 1 for further information.

We expressed the temperature-sensitive Dynamin mutant UAS-*Shibire^TS^* (Shi) at the level of KCs, trained animals at the restrictive temperature, and tested for memory performance at permissive temperature 30 minutes later. These manipulations allowed us to interfere with the synaptic vesicle exo-endocycle specifically during conditioning, while reinstating neurotransmission afterwards. Consistent with previous reports^28,30,31^, a slight drop in aversive memory performance (Fig. 1a) was not statistically different from their genetic controls, and also observable in the permissive temperature controls (see Supplementary Fig. 1a). In contrast, memories were completely abolished following block of KC output during appetitive training (Fig. 1b, Supplementary Fig. 1b).

We next asked whether the requirement for neurotransmission during appetitive learning was specific to the KC output synapse. To do so, we took an analogous approach, this time blocking neurotransmission from downstream M4/6 (MBON-γ5β’2a, MBON-β’2mp, MBON-β2β’2a and MBON-β’2mp bilateral) MBONs during appetitive training. We focused on the M4/6 set of MBONs as blocking these during memory retrieval crucially interferes with appetitive memory expression, while, on a physiological level, memory-related plasticity is observable^5,7,13^. When blocking M4/6 during appetitive training, but not retrieval, memory scores were similar to those of control groups (Fig. 1c), suggesting that the sites of plasticity are likely to be the KC to MBON synapse in general, with one major site specifically being the connections between KCs and M4/6 MBONs.

Thus, our experiments suggest that neurotransmitter release from KCs during training is required for the formation of appetitive memories, but is less crucial for the formation of aversive memories.

### The α5 nAChR subunit is required for induction, and α2 for expression of appetitive memories

Requirement for presynaptic neurotransmitter release alone does not necessarily mean that postsynaptic plasticity is involved in appetitive memory formation. To address a putative role for postsynaptic sensitivity in memory formation, we next interfered with the postsynaptic receptor composition. Given that KCs are cholinergic, we screened for memory requirement of all nicotinic α-subunits at the level of the M4/6 MBONs (Fig. 2) using previously published^4,51^ genetically-targeted RNAi. We concentrated on the nAChR α subunits, as they are crucial components for all possible heteromeric or homomeric receptor pentamers^36^.

**Figure 2:**
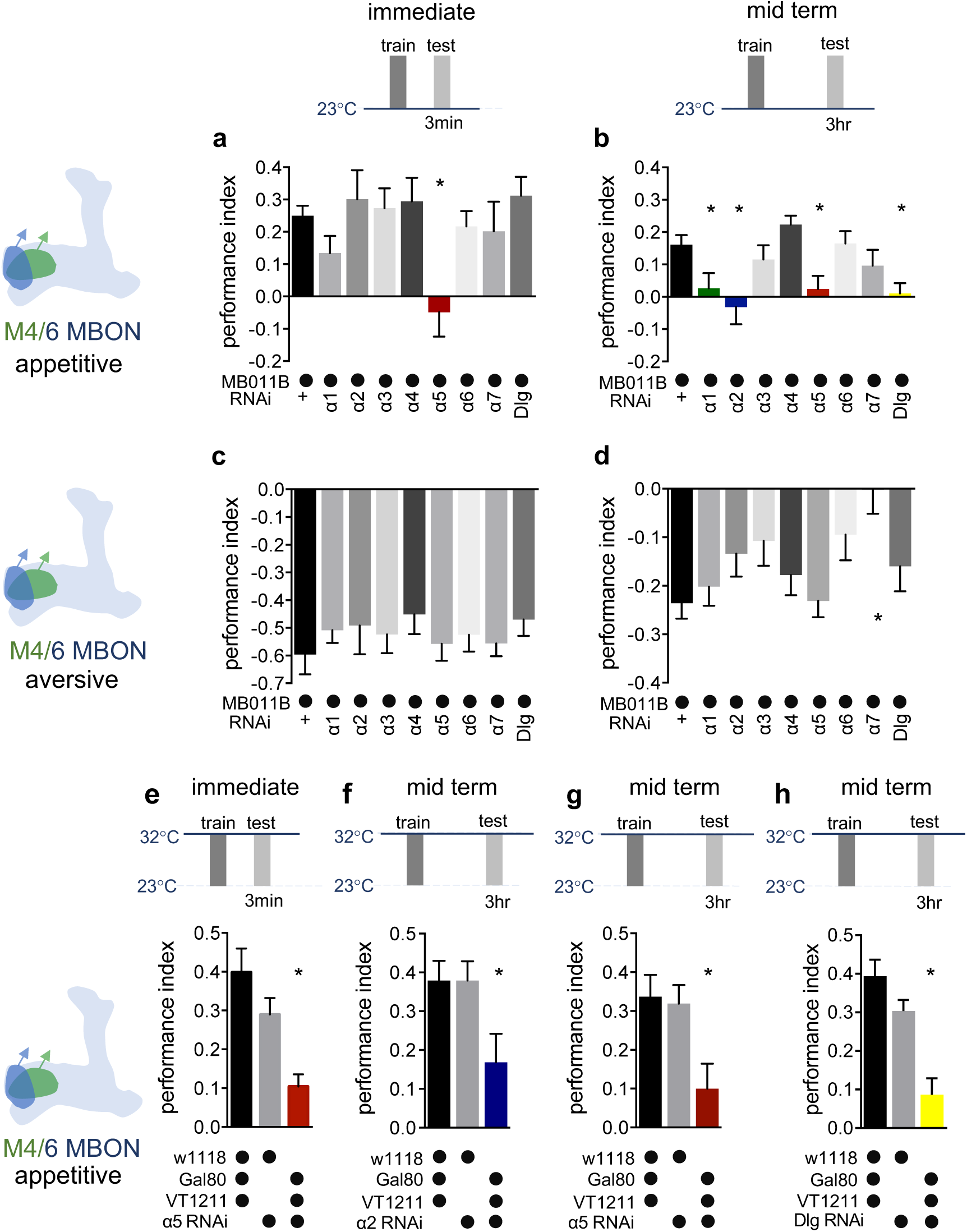
Specific nAChR α subunits are needed for specific memories in M4/6 neurons. **a)** Immediate appetitive memories are impaired following RNAi knock-down of the α5 nAChR subunit in M4/6 MBONs (driver line MB011B). Bar graphs: mean ± SEM; n = 8 – 13, for controls: n = 20; one-way ANOVA followed by Dunnett’s test (p < 0.05), * = p < 0.05. Note: data depicted correspond to initial screen, please see Supplementary Fig. 2i for alternate display including all genetic controls. **b)** RNAi knock-down of the α1, α2, α5 nAChR subunits or Dlg in M4/6 MBONs (driver line MB011B) impair 3-hour appetitive memories. Bar graphs: mean ± SEM; n = 12 – 26, for controls: n = 38; Kruskal-Wallis followed by Dunn’s test (p < 0.05), * = p < 0.05. Note: data depicted correspond to initial screen, please see Supplementary Fig. 2j-m for alternate display including all genetic controls. **c)** Immediate aversive learning is not impaired by RNAi knock-down of any subunit in M4/6 MBONs (driver line MB011B). Bar graphs: mean ± SEM; n = 6 – 8, for controls: n = 12; Kruskal-Wallis followed by Dunn’s test (p > 0.05). **d)** 3-hour aversive memory is not affected by knock-down of α subunits with the exception of α7 (driver line MB011B). Bar graphs: mean ± SEM; n = 21 – 32, for controls: n = 61. Kruskal-Wallis followed by Dunn’s test (p < 0.05), * = p < 0.05. **e)** RNAi knock-down of the α5 subunit in M4/6 MBONs (driver line VT1211-Gal4) is suppressed during development using Gal80^ts^. 3-5 days before the experiment RNAi-knock-down was induced. Immediate memory is significantly impaired. Bar graphs: mean ± SEM; n = 6 – 7; Kruskal-Wallis followed by Dunn’s test (p < 0.05), * = p < 0.05. **f)** RNAi knock-down of the α2 subunit in M4/6 MBONs (driver line VT1211-Gal4) is suppressed during development using Gal80^ts^. 3-5 days before the experiment RNAi knock-down was induced. 3-hour memories are significantly impaired. Bar graphs: mean ± SEM; n = 16 – 17; one-way ANOVA followed by Tukey’s test (p < 0.05), * = p < 0.05. **g)** RNAi knock-down of the α5 subunit in M4/6 MBONs (driver line VT1211-Gal4) is suppressed during development using Gal80^ts^. 3-5 days before the experiment RNAi knock-down was induced. 3-hour memories are significantly impaired. Bar graphs: mean ± SEM; n = 25 – 27; one-way ANOVA followed by Tukey’s test (p < 0.05), * = p < 0.05. **h)** RNAi knock-down of Dlg in M4/6 MBONs (driver line VT1211-Gal4) is suppressed during development using Gal80^ts^. 3-5 days before the experiment RNAi knock-down was induced. 3-hour memories are significantly impaired. Bar graphs: mean ± SEM; n = 8 – 11; one-way ANOVA followed by Tukey’s test (p < 0.05), * = p < 0.05. Also see Supplementary Fig. 2 for further information.

When flies were tested for immediate appetitive memory, only knock-down of the α5 subunit produced performance that was statistically different from the controls (Fig. 2a, Supplementary Fig. 2a,i). Testing 3-hour appetitive memory performance revealed a significant memory impairment in flies with α5, α1 and α2 knock-down (Fig. 2b, Supplementary Fig. 2b,j-l). While α5 subunits can form homomeric channels^38^, α1 and α2 can partake in heteromeric channels together^37^. We therefore concentrated on the α5 and α2 nAChR subunits in subsequent analyses.

To exclude developmental contributions to the observed memory defects, we repeated the immediate and 3-hour appetitive memory experiments for α5 as well as the 3-hour appetitive memory experiments for α2 knock-down animals, while suppressing RNAi expression using the temperature-sensitive Gal4 repressor Gal80^ts^ during development, up until 3-5 days before memory testing. Memory impairments were confirmed in all cases (Fig. 2e-g), but not detected in temperature controls (Supplementary Fig. 2e-g).

We also tested aversive immediate and 3-hour memory using the same genetic settings (Fig. 2c,d, Supplementary Fig. 2c,d). None of the knock-downs differed significantly from controls, with the exception of α7 at the 3-hour time point. As, comparable to vertebrate systems, α7 also plays a significant role at presynaptic neurites^39^, we did not follow up on this observation in this study.

As M4/6 output is also required for appropriate aversive memory expression^7,11^, α2 and α5 knock-down not impacting aversive memory performance suggested that the observed appetitive memory impairments were not simply a consequence of lost postsynaptic sensitivity to acetylcholine. To further corroborate this, we turned to a brain explant preparation and applied acetylcholine focally to the dendrites of M4/6 neurons that expressed the calcium indicator GCamp6f, for both control and knock-down settings, in the presence of the blocker of voltage-gated sodium channels TTX^4,40^. Dendritic calcium transients were comparable between all groups (Supplementary Fig. 3e). We also observed presynaptic sensitivity in all genotypes (*not shown*) after applying acetylcholine to the presynaptic MBON boutons, making presynaptic deficits following α2 or α5 knock-down unlikely.

Therefore, we conclude that, at the level of M4/6 neurons, immediate and 3-hour appetitive memories are affected by knock-down of the α5 subunit, whereas 3-hour memories also require the presence of α1- and α2-bearing receptors in addition. The observed temporal profile of requirement for memory of α1- and α2-bearing receptors relative to those incorporating the α5 subunit, could potentially point to a temporal sequence of receptor function during initial memory formation and subsequent memory expression.

### The postsynaptic scaffold Dlg is required for 3-hour appetitive memory

At mammalian glutamatergic synapses postsynaptic receptor-mediated changes in synaptic weight rely on receptor stabilization or destabilization that can be mediated via scaffolding molecules. One such scaffold, PSD-95, that is mostly involved in AMPA receptor motility, is conserved at *Drosophila* synapses. The orthologue Dlg^41,42^ is expressed throughout the brain, with mushroom body compartment-specific enrichment noted previously^43^ (also compare Fig. 4a,b). We investigated appetitive and aversive memory performance following M4/6-specific knock-down of Dlg (Fig. 2a-d, Supplementary Fig. 2a-d). Performance scores comparable to controls were found for both immediate appetitive and aversive memories (Fig. 2a,c, Supplementary Fig. 2). Dlg knock-down, however, specifically abolished 3-hour appetitive memory performance (Fig. 2b,d, Supplementary Fig. S2b), while Gal80^ts^ experiments excluded a developmental defect (Fig. 2h, Supplementary Fig. 2h). The temporal profile of Dlg requirement therefore closely matched that of α2 nAChR subunits.

### Bypassing the presynapse: induction of persistent associative plasticity in the postsynaptic compartment

Recent ultrastructural data has revealed direct synaptic connections between dopaminergic neurons and MBONs^23,44^, giving rise to a motif that could allow for postsynaptic plasticity induction (see schematic in Fig. 3a). In order to directly test whether postsynaptic plasticity could take place at the level of MBONs, we next conducted neurophysiological proof-of-principle experiments.

**Figure 3:**
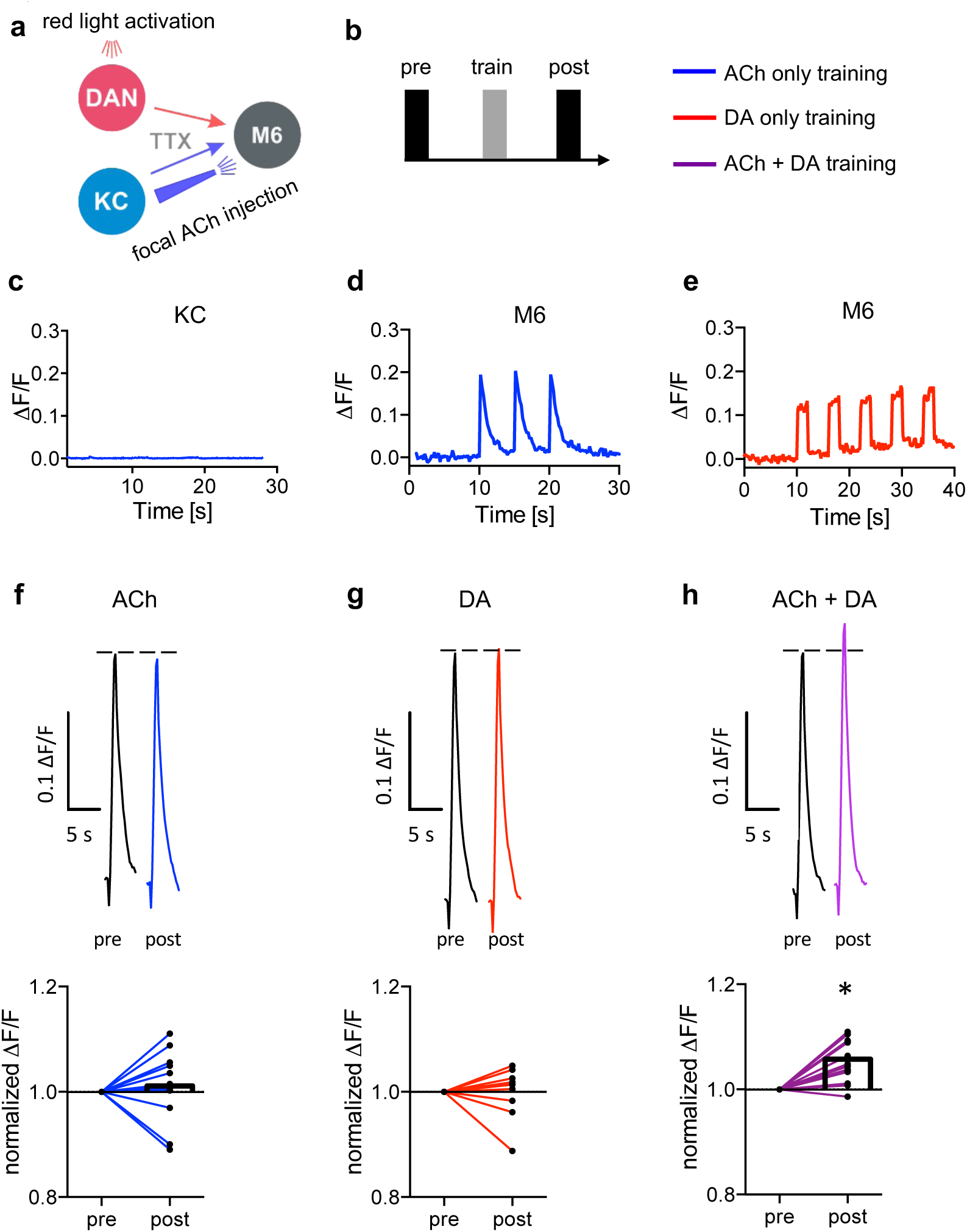
Induction of postsynaptic plasticity bypassing the presynapses. **a)** Connectivity scheme of MB output synapses. Cholinergic KCs and dopaminergic neurons are presynaptic to M6 MBONs. Only connections relevant for this protocol are shown for simplicity. Red light pulses trigger release of dopamine (DA) from dopaminergic neurons (R58E02-LexA > lexAop-CsChrimson^tdTomato^), while KC input is circumvented and mimicked by focal acetylcholine (ACh; 0.1 mM) injections to M6 dendrites in an explant brain preparation. Postsynaptic responses at the level of M6 are measured using GCaMP6f (MB011B > UAS-GCaMP6f). TTX in the bath suppresses feedback signaling and overall network activity within the circuit. **b)** Training scheme (top). Baseline responses to ACh application were initially established (pre). Subsequent training protocols consist of either pairing ACh application with simultaneous activation of dopaminergic neurons (purple connection lines), activation of dopaminergic neurons (‘red light only, red connection lines), or ACh only (blue, connection lines). This is followed by a test trial (post) through ACh application. **c)** Averaged traces of axonal KC calcium changes induced by focal ACh injections. No apparent transients are observable; n = 7. Line is mean ± SEM. **d)** Sample trace of dendritic M6 calcium changes induced by focal ACh injections. **e)** Sample trace of dendritic M6 calcium changes induced by red-light pulses. **f-h)** Above: sample calcium traces in response to ACh injections recorded from M6 dendrites pre (black traces) and post (colored traces) training. Below: peak quantification. **f)** Changes in acetylcholine-evoked calcium transients; comparison of mean peaks pre- and post ‘ACh only’ training. Before-after plots and bar graphs (mean); n = 13, ratio paired t-test. **g)** Changes in acetylcholine-evoked calcium transients; comparison of mean peaks pre- and post ‘red light only’ training. Before-after plots and bar graphs (mean); n = 10, ratio paired t-test. **h)** Changes in acetylcholine-evoked calcium transients; comparison of mean peaks pre- and post ‘paired’ training. Before-after plots and bar graphs (mean); n = 18, ratio paired t-test, * = p < 0.05. Also see Supplementary Fig. 3 for further information

To minimize plasticity induced by acute sensory experiences or general network activity, we used an explant brain preparation bathed in TTX from flies expressing the red light-activatable channelrhodopsin CsChrimson in a subset of dopaminergic neurons (PAM neurons; R58E02-LexA) and the calcium indicator GCaMP6f in M4/6 MBONs.

While dopamine release was controlled by red light flashes, neurotransmitter release from KCs was mimicked by focal pressure ejection of acetylcholine to the dendrites of the M6 (MBON-γ5β’2a) MBON (M6 was chosen for technical reasons, as these neurons are most accessible for the used imaging technique). We verified that KC presynapses do not respond to acetylcholine application^4^, using both calcium imaging and imaging of synaptic vesicle exocytosis at the level of KC axons (Fig. 3c and Supplementary Fig. 3f,g). The observed absence of KC activation, with acetylcholine being applied from an external source (Fig. 3a), minimized noise attributable to possible presynaptic contributions.

Our protocols consisted of training phases where we differentiated between temporal pairing of acetylcholine and optogenetic activation of dopaminergic neurons (‘paired’, Fig. 3b,h), dopamine only (‘red-light only’, Fig. 3b,e,g), or ‘acetylcholine only’ (Fig. 3b,d,f). Acetylcholine application preceded (pre) and followed each training step (post) to establish baseline responses and to assess synaptic weights following training (‘testing’). We found that test responses were significantly elevated following the paired condition (Fig. 3h). This potentiation was not observed when testing after acetylcholine only or dopamine only training (Fig. 3f,g). Importantly, we also did not observe any changes when pairing acetylcholine application with red light in non-CsChrimson-expressing controls (Supplementary Fig. 3c,d).

Because we are using global acetylcholine application instead of sparse activation of single synapses, these experiments likely do not reflect *in vivo* physiological settings^7^. However, our proof of principle experiments demonstrate that postsynaptic sensitivity can change independently of the presynapse. Indeed, the observed postsynaptic potentiation was not observed, when knocking-down α2 in M4/6 (Supplementary Fig. 3h), consistent with postsynaptic plasticity being linked to the requirement of nicotinic receptors in memory storage.

### Non-uniform distribution of nAChR α-subunits throughout MB compartments

Our data so far are suggestive of α2-containing nicotinic receptors being involved in appetitive memory storage. To test whether receptor levels were interdependent, we made use of a newly established CRISPR-based genomic collection of GFP-tagged endogenous nAChR subunits (Woitkuhn, Pribbenow, Matkovic, Sigrist and Owald, *unpublished*) covering all α subunits (with the exception of α3) under control of their endogenous promoter, allowing for analyses of receptor distribution and dynamics in a dense neuropile *in situ*.

We first characterized receptor subunit signals throughout the 15 MB compartments, several of which have been shown to be involved in the encoding of specific memories. We found a non-uniform distribution (Fig. 4a,b, Supplementary Fig. 4) that was unique for each subunit, indicating considerable heterogeneity of receptor composition.

**Figure 4:**
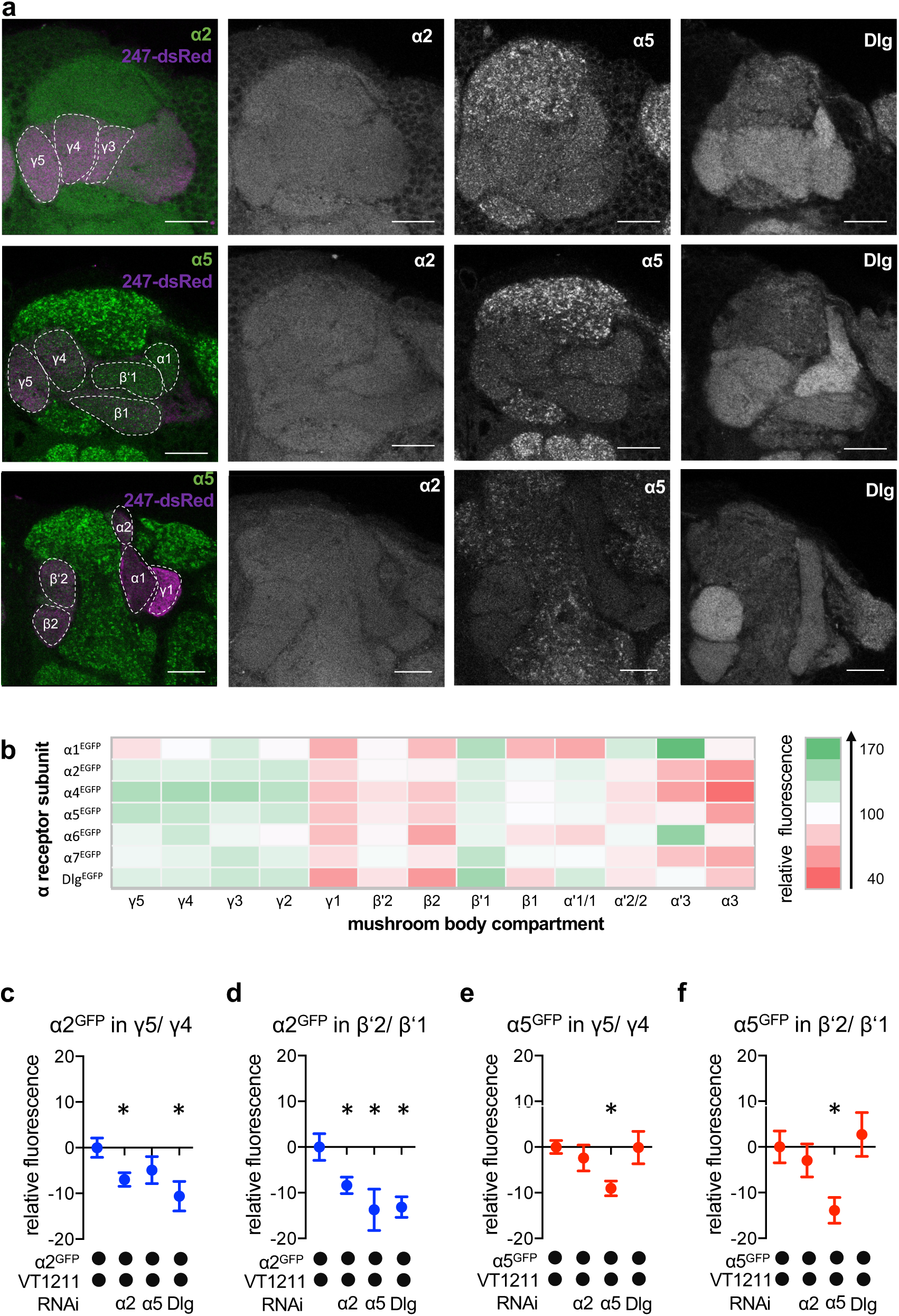
nAChR α subunit localization throughout the MB: MBON-specific RNAi alters subunit distribution. **a)** Representative images of the GFP-tagged nAChR subunits α2 and α5 as well as Dlg. Individual images displayed here are taken from different animals. For other subunits see Supplementary Fig. 4. Scale-Bar: 20 μm. Left: merge of α subunit signal (green) with MB compartments marked with 247-dsRed (magenta). Compartments are indicated by dashed-lines. Top row: γ compartments; middle row: α, β’, β and γ compartments, bottom row: α’, α and β compartments. **b)** Quantification of all GFP-tagged α receptors (except for the α3 subunit). GFP signals for the indicated MB compartments are relative to the mean intensity of the GFP signal of the complete MB; n = 7 – 18. **c)** Knock-down of Dlg or α2 in M4/6 neurons (driver line VT1211-Gal4) significantly reduces the α2^GFP^ fluorescence within the γ5 compartment (relative to unmanipulated γ4). Bar graph: normalized mean ± SEM; n = 8 – 20; Kruskal-Wallis followed by Dunn’s test (p < 0.05), * = p < 0.05. **d)** Knock-down of either the α2 or the α5 nAChR subunit or Dlg in M4/6 neurons (driver line VT1211-Gal4) decreases the relative fluorescence signal of α2^GFP^ in the β’2 compartment (relative to unmanipulated β’1). Bar graph: normalized mean ± SEM; n = 9 – 20; one-way ANOVA followed by Dunnett’s test (p < 0.05), * = p < 0.05. **e)** Knock-down of α5 in M4/6 neurons (driver line VT1211-Gal4) decreases the α5^GFP^ signal in the γ5 compartment (relative to unmanipulated γ4). Bar graph: normalized mean ± SEM; n = 9 –28; Kruskal-Wallis followed by Dunn’s test (p < 0.05), * = p < 0.05. **f)** α5^GFP^ fluorescence is significantly decreased in the β’2 compartment (relative to unmanipulated β’1) after knock-down of α5 in M4/6 neurons (driver line VT1211-Gal4). Bar graph: normalized mean ± SEM; n = 10 – 28; Kruskal-Wallis followed by Dunn’s test (p < 0.05), * = p < 0.05. Also see Supplementary Fig. 4 for further information.

α5, which is required for immediate and 3-hour appetitive memories, was abundant throughout the γ lobe, including γ5 (innervated by M6) and slightly less at the level of β’2 (innervated by M4 and in parts by M6). α2 subunits, required for 3-hour appetitive memories, showed similarly high relative abundance in β’2 and γ5 (Fig. 4a,b). Of note, these subunits were also detected in other MB output compartments, such as α’3, which harbors MBONs involved in non-associative familiarity learning^15^.

We next evaluated, whether the fluorescent signals of the α2 and α5 subunits (with α5 potentially functioning upstream of α2, Fig. 2) observed in the β’2 and γ5 compartments were derived from receptors within the dendritic processes of M4/6. To do so, we performed cell-specific knock-down experiments using VT1211-Gal4 and quantified the relative fluorescent signal of the knock-down compartment relative to the neighboring unmanipulated compartments (Fig. 4c-f).

Knock-down of the α5 nAChR subunit reduced the relative α5^GFP^ signal specifically and significantly in γ5 and β’2 (Fig. 4e,f). α5 abundance was, however, unaltered when knocking-down α2 or Dlg, which is in line with α5 functioning as a possible trigger for plasticity processes.

Likewise, confirming that the observed signal was derived from M4/6 MBON dendrites, α2 knock-down reduced relative α2^GFP^ levels significantly throughout the β’2 and γ5 compartments (Fig. 4c,d). Strikingly, we also observed reduced α2 nAChR subunit levels following α5 subunit knock-down in the β’2 compartment or Dlg knock-down in the β’2 and γ5 compartments (Fig. 4c,d), which would be in line with a Dlg-dependent sequential requirement of receptor subunits during memory formation (also compare behavioral data in Fig. 2).

Our data therefore are consistent with a role of α5 nAChR subunits and Dlg functioning upstream of α2 subunit-positive receptors, at least within the β’2 compartment.

### nAChR subunits shape synaptic MB output properties

We next focused on implications of α2 subunit knock-down on postsynaptic function of M4/6 MBONs. Axonal calcium transients had previously been shown to be decreased following knock-down of α subunits^4^. However, both increased or decreased postsynaptic drive could lead to changed dendritic integration properties underlying reduced signal propagation^45^.

We expressed GCaMP6f in M4/6 MBONs (Fig. 5a), and exposed the flies repeatedly to alternating puffs of the odors octanol (OCT) and MCH (Supplementary Fig. 5a,b). We focused our experiments on the β’2 compartment (Fig. 5), as this is innervated by M4 MBONs that show input-specific plasticity following appetitive learning^7^. Initial dendritic odor responses were not statistically different between α2 subunit knock-down and controls (Fig. 5b,c), while initial odor-evoked dendritic calcium transients were significantly elevated following knock-down of α5 (Fig. 5b,c). Importantly, while control animals showed a relative facilitation in odor-specific calcium transients after several exposures to OCT, we did not detect this in α2 knock-down animals (Fig. 5d,e). Odor responses following α5 knock-down, however, clearly depressed after multiple odor exposures (Fig. 5f), indicating that lack of α5 can lead to pre-potentiated synaptic transmission, while α2 nAChR subunit knock-down interfered with odor-evoked baseline transmission properties to a lesser extent. We did not observe any changes in calcium signals at the level of the corresponding KC axons, further supporting that the observed plasticity was of postsynaptic origin (Supplementary Fig. 5h,i).

**Figure 5:**
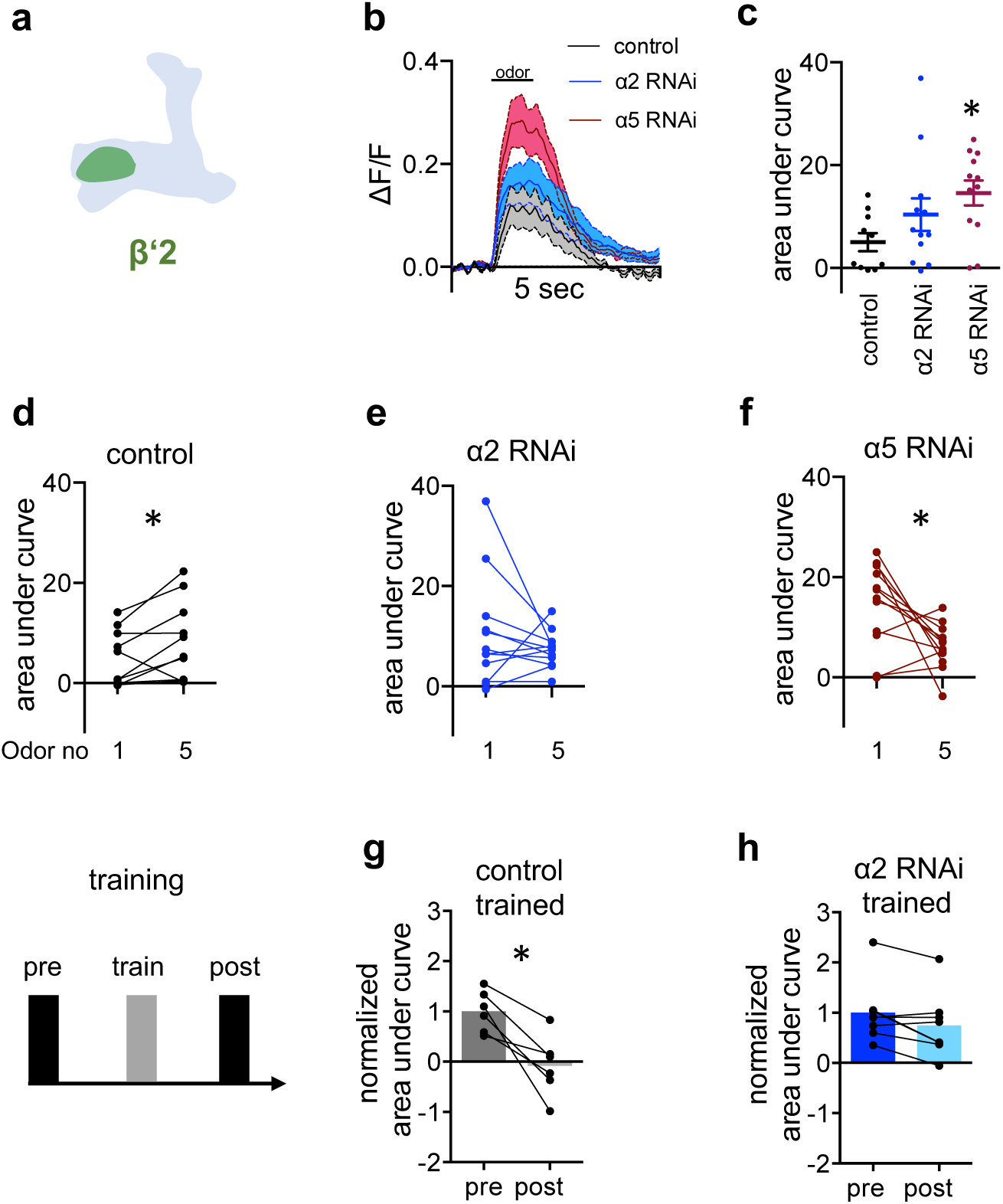
α2 is required for learning-associated plasticity *in vivo*. **a)** Scheme indicating the dendritic imaging area **(b-f)** at the level of the β’2 compartment. **b)** Averaged traces of GCaMP6f responses to OCT from control (black), α2 subunit RNAi (blue) and α5 subunit RNAi (red; driven in M4/6 respectively; VT1211-Gal4 as driver line) flies. Solid traces are mean, shaded areas SEM; n = 10 – 12. Odor exposure is indicated by bar. **c)** Area under curve quantifications of averaged odor responses show significantly elevated odor responses to OCT following α5 knock-down in M4/6 neurons (driver line VT1211-Gal4). Mean ± SEM; n = 10 – 12; Kruskal-Wallis followed by Dunn’s test (p < 0.05); * = p < 0.05. **d)** Control flies show a significant increase between the first and the fifth response to OCT. Before-after plots; n = 10; paired t-test; * = p < 0.05. **e)** α2 RNAi flies show no difference between the first and fifth odor response to OCT. nAChR subunit RNAi is driven in M4/6 neurons (driver line VT1211-Gal4). Before-after plots; n = 12; Wilcoxon matched-pairs signed rank test. **f)** α5 RNAi flies show a significant decrease in calcium transients over the course of consecutive odor exposures. nAChR subunit RNAi is driven in M4/6 neurons (driver line VT1211-Gal4). Before-after plots; n = 12; paired t-test; * = p < 0.05. **g)** Control flies show a significant decrease in GCaMP6f responses to MCH (trained odor) following absolute training (axonal imaging). Before-after plots and bar graphs (mean); n = 6; Wilcoxon matched-pairs signed rank test; * = p < 0.05. **h)** α2 RNAi flies show no significant decrease in GCaMP6f response to MCH (trained odor) following absolute training (driver line VT1211-Gal4, axonal imaging). Before-after plots and bar graphs (mean); n = 8; Wilcoxon matched-pairs signed rank test. Also see Supplementary Fig. 5 for further information.

### *The α2* nAChR subunit is required for the formation of appetitive memory traces *in vivo*

Of note, the observed facilitation in M4/6 of controls was not apparent when using repeated application of MCH, indicating that parameters underlying the observed non-associative plasticity are odor-dependent (Supplementary Fig. 5e). However, the only slight effects on base-line MCH-induced responses observed for α2 knock-down animals, allowed us to next investigate the role of α2 during *in vivo* associative appetitive learning.

We performed *in vivo* training under the microscope experiments using an absolute paradigm, pairing odor (MCH) exposure with *ad libitum* sugar feeding during training. Comparing odor responses at the level of M4 before and after training revealed a marked depression for control animals in line with previous observations^7,17^ (Fig. 5g). Importantly, after MBON-specific knock-down of α2, this training-induced plasticity was not observed (Fig. 5h).

Together, our data point towards a mechanism, where nicotinic receptor subunits shape synaptic properties (Fig. 5), with α2 as a postsynaptic substrate underlying appetitive training-induced plasticity processes.

### *In vivo* imaging of postsynaptic receptor plasticity reveals altered α2 dynamics

Structural changes at the level of the receptor composition are hallmarks of postsynaptic plasticity expression in vertebrates. Typically, rearrangements can be measured by altered dynamics (or motility) of receptors that can reflect incorporation or removal of receptor complexes. We next sought to test whether dynamic receptor behavior could serve as a structural correlate of cholinergic postsynaptic memory trace expression. To do so, we turned to *in vivo* imaging experiments of the endogenously tagged α2 subunit (Fig. 6, Supplementary Fig. 6). Flies were either tethered under the two (Fig. 6) or the single photon confocal (Fig. 7) microscope and subjected to training protocols. Endogenous *in situ* receptor dynamics at the level of the β’2 compartment of the MB were estimated as fluorescence recovery after photobleaching (FRAP), allowing us to also pick up small increases in signal (Fig. 6a-c).

**Figure 6:**
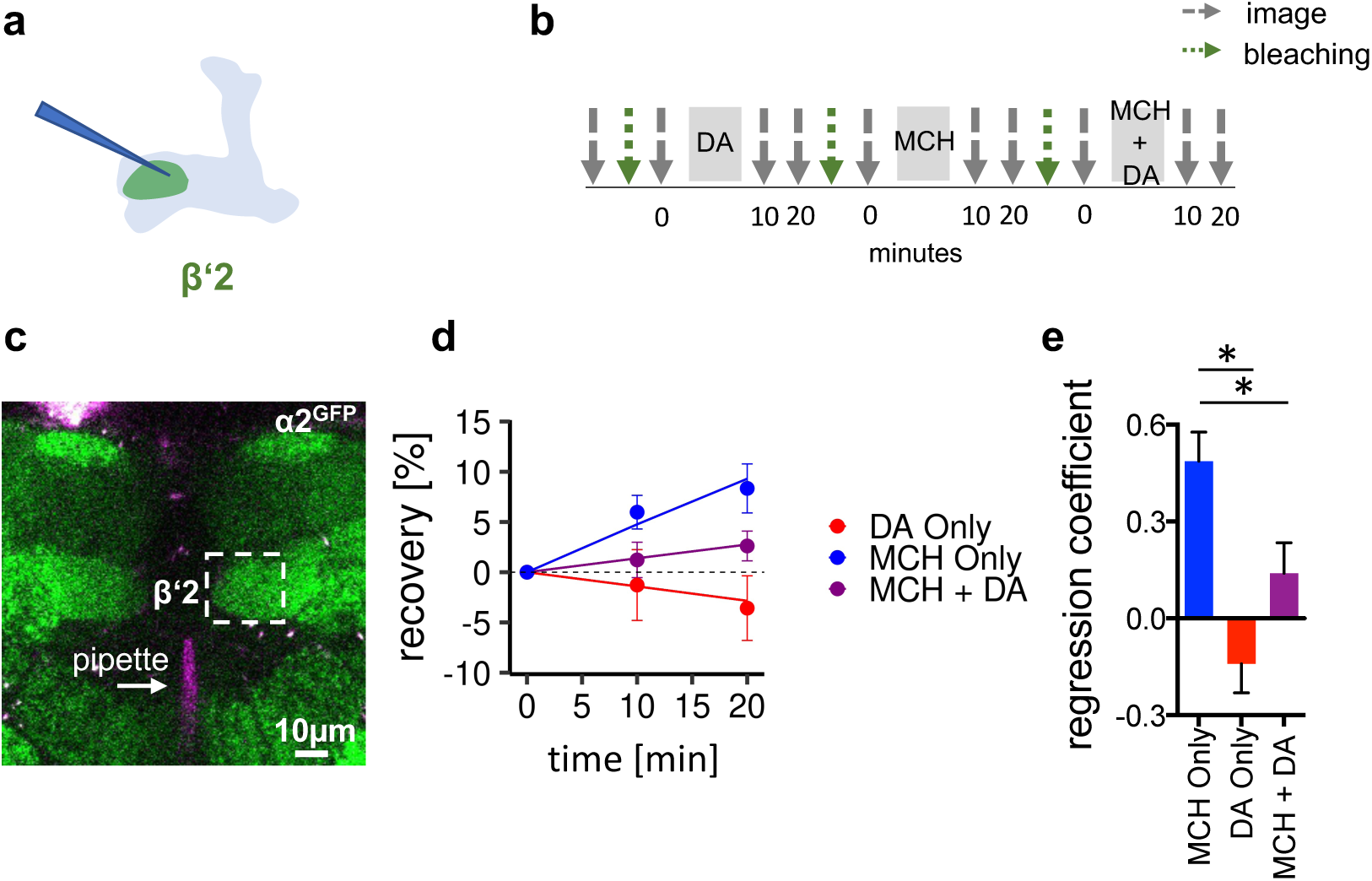
α2 nAChR subunits dynamically rearrange. **a)** Scheme of site of dopamine injection during fluorescence recovery after photobleaching (FRAP) experiments at the level of the KC-MBON synapses of the β’2 compartment. **b)** FRAP experimental protocol. After bleaching the baseline picture was taken followed by dopamine injection, odor presentation and odor presentation simultaneously with dopamine injection in the same fly. Fluorescence recovery was monitored at the 10- and 20-minute time points. **c)** Example image of α2^GFP^ expression; white dashed box shows the β’2 output zone; dopamine injection pipette (with Texas Red) is labelled in magenta. Scale bar: 10 µM. **d)** Regression of fluorescence recovery after photobleaching following MCH exposure (blue line), MCH exposure simultaneously with dopamine (DA) injection (purple line), dopamine injection alone (red line). **e)** Regression coefficient. Bar graphs: regression coefficients of recovery kinetics ± standard error of regression; n = 9 – 10, linear mixed effects model followed by pairwise comparison from estimated marginal trends. * = p < 0.05. Also see Supplementary Fig. 6 for further experiments.

**Figure 7:**
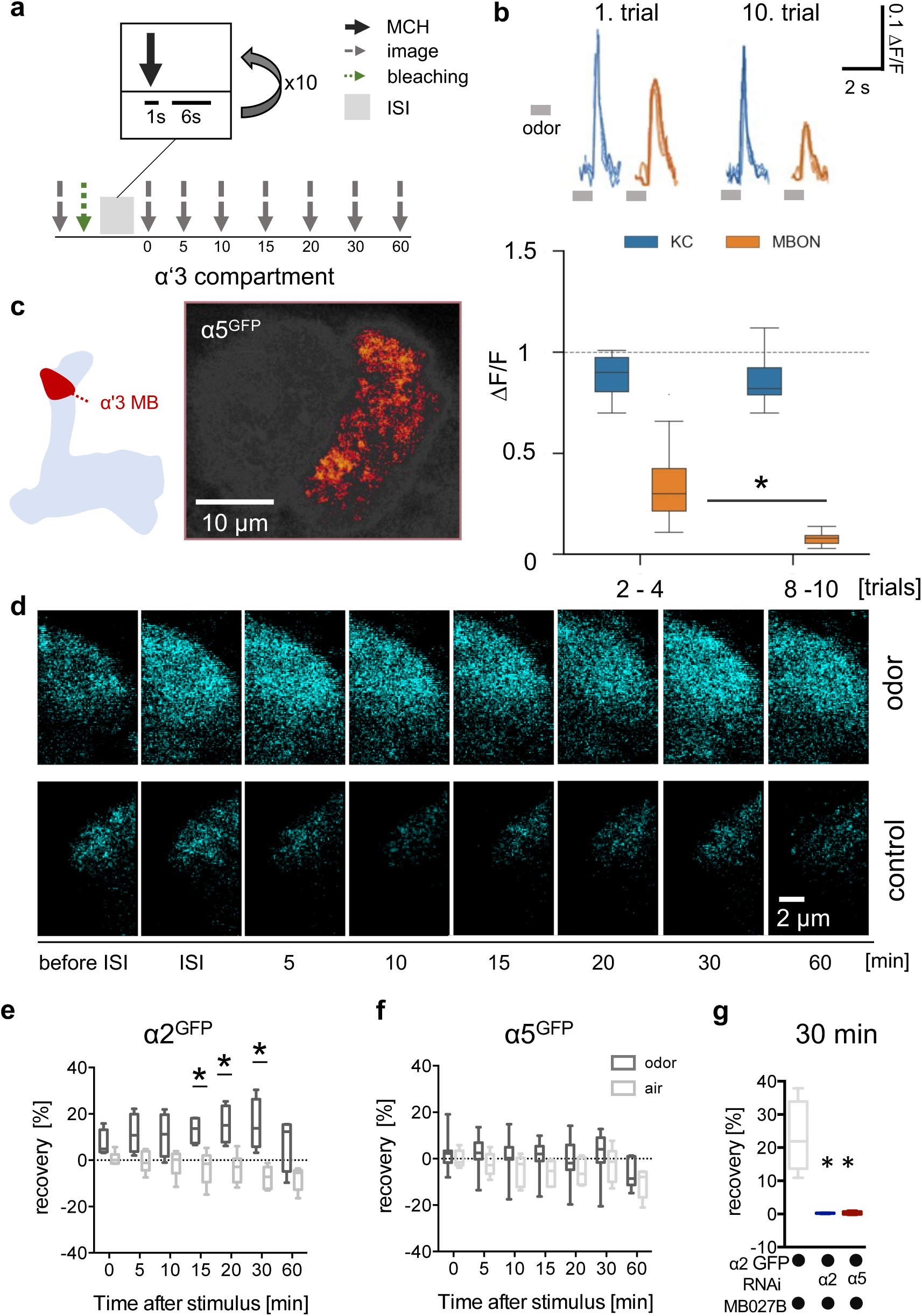
Non-associative plasticity alters postsynaptic α2 subunit receptor dynamics. **a)** Training scheme indicating odor application, bleaching and imaging time points. MCH was given 10 times for 1 second with a pause of 6 seconds in-between. Images were taken after training in absence of odor immediately afterwards and 5, 10, 15, 20, 30 and 60 minutes later. **b)** Top: calcium peaks in response to odor stimuli of presynaptic KCs (MB369B as driver line) and adjacent postsynaptic MBONs (driver line: MB027B). Individual calcium responses to trials 1 and 10 for MBONs (orange lines) and KCs (blue lines). Bottom: Averaged calcium responses to odor stimuli of presynaptic KCs and postsynaptic MBONs of trials 2 - 4 and 8 - 10 respectively. Responses decrease at the level of MBONs but not at the level of KCs over ten trials. Box plots are median and 75 % quartiles; n = 15; Kruskal-Wallis followed by Dunn’s test (p < 0.05), * = p < 0.05. **c)** Scheme of α’3 compartment analyzed and representative α5^GFP^ fluorescent image (smoothed). Scale bar: 10 μm. **d)** Example images of α2^GFP^ FRAP experiment at the level of the α’3 compartment at specific time points before and after training. Top row, after training; bottom row: control settings. Scale bar: 2 μm. **e)** FRAP of α2^GFP^ nAChR subunit in the α’3 compartment after odor presentation. α2^GFP^ shows significant recovery following odor training compared to the controls. Recovery rate is normalized to the baseline recorded after selective bleaching of the α’3 MB compartment. Box plots are median and 75 % quartiles; n = 4 – 6; multiple t - tests with Sidak-Bonferroni correction, * = p < 0.05. **f)** FRAP of α5^GFP^ subunit in the α’3 compartment after odor presentation. α5^GFP^ did not show significant recovery compared to the controls. Recovery rate is normalized to the baseline recorded after selective bleaching of α’3 MB compartment. Box plots are median and 75 % quartiles; n = 5 – 7, multiple t - tests with Sidak-Bonferroni correction. **g)** FRAP of α2^GFP^ nAChR subunit in the α’3 compartment after odor presentation and knockdown of either the α2 or α5 subunit in the α’3 MBON (driver line MB027B). α2^GFP^ shows no recovery 30 min after odor training in α2 or α5 knockdown animals compared to the controls. Recovery rate is normalized to the baseline recorded after selective bleaching of the α’3 MB compartment. Box plots are minimum value to maximum value; n = 4 – 5; Kruskal-Wallis followed by Dunn’s test (p < 0.05), * = p < 0.05. Also see Supplementary Fig. 7 for further information.

We conducted artificial appetitive training protocols by exposing the tethered animals to odor with or without simultaneous focal injection of dopamine to the β’2 compartment of the MB (Fig. 6a-c). We performed two different sets of protocols comparing recovery of α2^GFP^ (Fig. 6b, Supplementary Fig. 6b).

Following photobleaching, individual flies were exposed to consecutive focal injections of dopamine, odor, and odor paired with focal dopamine injection. MCH induced significantly increased fluorescence recovery when compared to dopamine injections only or odor paired with dopamine (Fig. 6d,e). Dopamine, therefore, does not induce plasticity on its own, and further, it suppresses odor-induced recovery when applied simultaneously with an odor. To rule out that recovery depended on the type of odor used, we also conducted similar experiments using OCT, this time testing all conditions in separate flies. After photobleaching, flies were either exposed to a focal injection of dopamine, an odor stimulus, or odor paired with focal dopamine injection. We only observed significant recovery in the odor only condition, whereas dopamine and odor paired with dopamine induced no significant recovery (Supplementary Fig. 6a-d).

Thus, our data indicate that pairing odor presentation with dopamine application stalls α2^GFP^ dynamics, potentially by either stabilizing the already present amount of receptor or hindering new incorporation of α2-containing receptors. Interestingly the opposite, increased receptor dynamics, is observed after odor exposure without reinforcer. Note that the absence of acute stimuli (constant air stream only), did not induce signal recovery, demonstrating that it is the presence of the odor that changes baseline receptor behavior (Supplementary Fig. 6e). Thus, stalling α2 dynamics can be correlated to depression of M4/6 MBON synapses^7,13^ following appetitive training (Fig. 5g,h).

### Familiarity learning alters postsynaptic receptor dynamics

We next asked whether postsynaptic plasticity expressed through α5 and α2 subunit interplay could underlie other forms of learning represented in the MBs. We turned to the α’3 compartment at the tip of the vertical MB lobe that has previously been shown to mediate odor familiarity learning. This form of learning allows the animal to adapt its behavioral responses to new odors and, importantly, permits for assaying direct odor-related plasticity. Importantly, this compartment follows different plasticity rules, because the odor serves as both the conditioned (activating KCs) and unconditioned stimulus (activating corresponding dopaminergic neurons)^15^. While allowing us to test whether the so far uncovered principles could also be utilized in a different context, it also provides a less complex test bed to further investigate whether α5 functions upstream of α2 dynamics.

Confirming previous observations^15^, a repeated odor application paradigm (Fig. 7a) led to the depression of postsynaptic calcium transients at the level of the α’3 MBONs (Fig. 7b). Importantly, we did not detect a corresponding depression on the presynaptic side when imaging arbors of a sparse α’β’ KC driver line within α’3 (Fig. 7b), further indicating that memories were predominantly stored postsynaptically in this compartment. We next performed *in vivo* FRAP experiments following familiarity learning paradigms. After odor training, we observed clear recovery rates of α2^GFP^ signals compared to the control group, however not of α5^GFP^ or Dlg^GFP^ (Fig. 7 c-f, Supplementary Fig. 7). Therefore, increased α2 subunit dynamics are triggered through training events and, at the level of the α’3 compartment, accompany postsynaptic depression of the MBONs.

To invariantly test whether the observed recovery was attributable to α2 expressed in α’3 MBONs, we trained animals while knocking-down α2 specifically in α’3 MBONs. In accordance with the observed signal recovery deriving from MBONs, no recovery was observed after α2 knock-down (Fig. 7g). Importantly, we did not observe α2^GFP^ recovery when performing specific α5 knock-down in α’3 MBONs (Fig. 7g), indicating a role of α5 upstream of α2 also in this compartment.

### α5 subunits govern induction and α2 subunits expression of non-associative familiarity learning

Finally, we tested whether interfering with α5 and α2 nAChR subunits at the level of α’3 MBONs would also impact familiarity learning behavior (Fig. 8). Flies were covered in dust and subjected to repeated odor exposures^15^. As expected^15^, control flies readily groomed to remove the dust, however typically stopped this action when detecting the novel odor (Fig. 8a-c). Over subsequent trials, control flies learned that this odor was familiar and stopped reacting to the stimulus, continuing grooming (Fig. 8a-c, Supplementary Fig. 8). Expressing RNAi to the α2 subunit at the level of the α’3 MBONs clearly impacted learning: flies learned with decreased efficacy and only after several trials (Fig. 8a-d). Strikingly, α5 RNAi-expressing flies failed to stop grooming even to the first stimulus. Indeed, they acted as if they had already learned that an odor was familiar (Fig. 8a-d).

**Figure 8:**
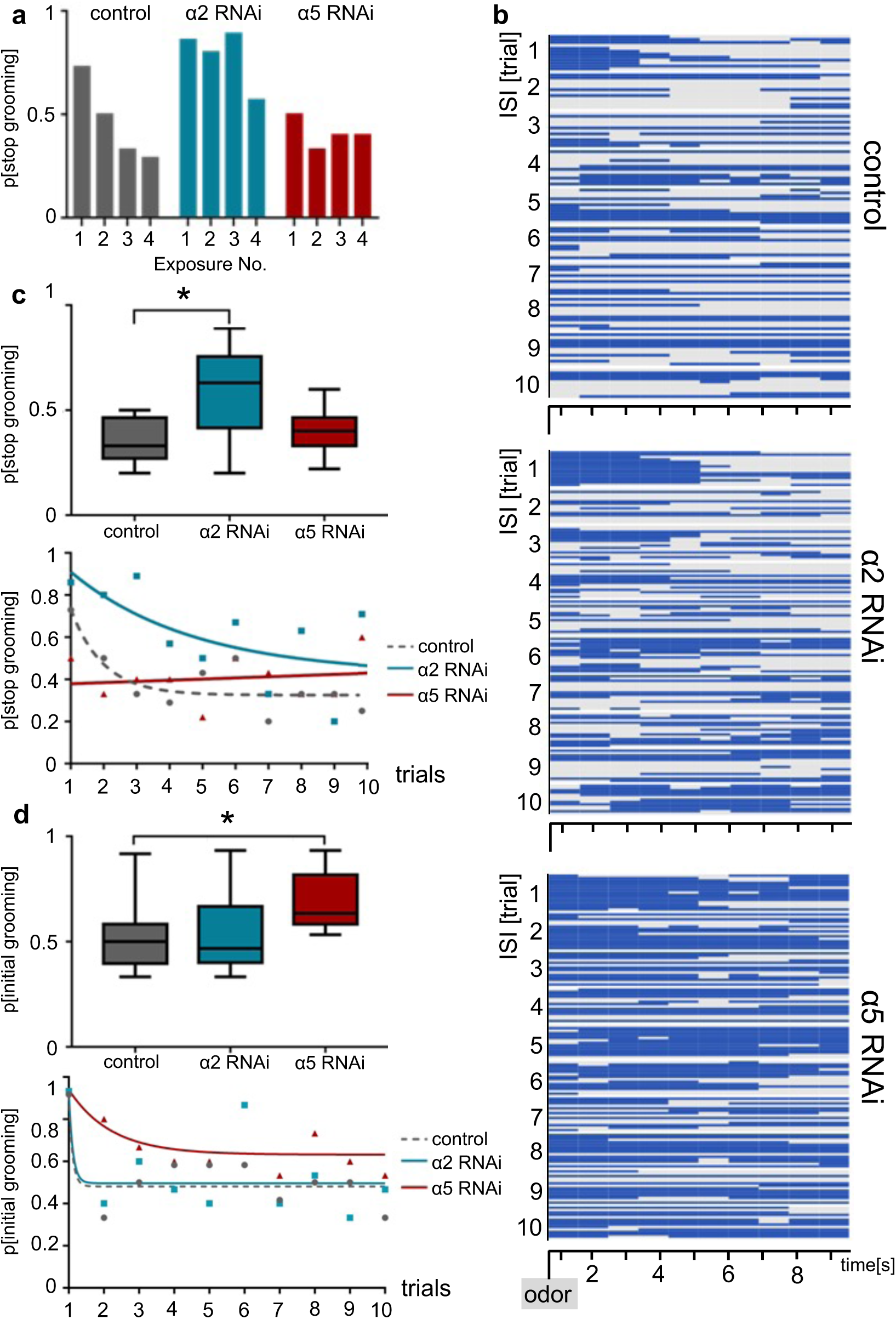
α2 and α5 nAChR subunits are required for non-associative familiarity learning at the level of α’3 MBONs. **a)** Knock-down of α nAChR subunits at the level of α’3 MBONs alters odor familiarity learning and the probability to stop grooming. α2 RNAi knock-down delays familiarity learning effects to novel odors. α5 RNAi knock-down flies do not show a novelty response at all. **b)** Grooming behavior response of dusted flies following the repeated presentations of a novel odor (MCH). Ethogram of grooming behavior (blue) during ten intervals of odor exposures. Horizontal lines in each trial correspond to a single experimental fly within a trial group. Not grooming (grey) flies can further be categorized between pausing and wandering (see Supplementary Figure 8). n = 15. **c)** Knock-down of α2 subunit in α’3 MBONs (driver line MB027B) impairs odor familiarity learning significantly by showing a higher probability to terminate grooming responses during the learning period. The learning period is defined as the odor exposure rounds following the first exposure). Bottom graph: non-linear representation of grooming flies over ten training trials. Note that α5 behavioral responses are best described by linear representation. Box plots are median and 75 % quartiles; n = 9, one-way ANOVA followed by Dunnett’s test (p < 0.05) * = p < 0.05. **d)** Knock-down of α5 subunits in α’3 MBONs (driver line MB027B) leads to an increased probability to start grooming earlier. ‘Grooming’ is defined here as constantly grooming during 2 and 3 seconds after odor delivery. Bottom graph, non-linear representation of grooming flies over ten training trials. Box plots are median and 75 % quartiles; n = 9, Kruskal-Wallis followed by Dunn’s test (p < 0.05), * = p < 0.05. Also see Supplementary Fig. 8 for further information.

Together, our data are in line with a model where α5 can induce memory formation, while lack of α5 leads to fully potentiated synapses. Subsequent expression of memory traces requires α2-containing receptors. Importantly, recovery accompanies synaptic depression at the level of the α’3 MBONs, while being suppressed by paired training in the β’2 compartment. Moreover, α2 appears to be involved in both depression and facilitation of synapses. Thus, synapses could bidirectionally utilize plasticity of the same receptor subunit for storing different types of information (Fig. 9).

**Figure 9:**
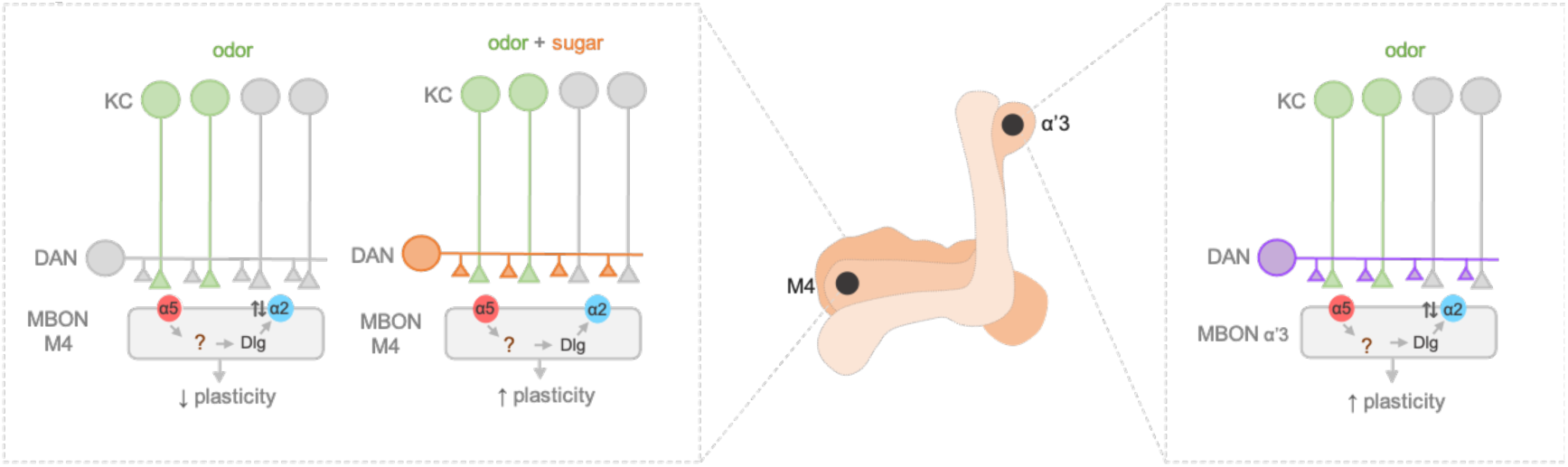
Model of postsynaptic plasticity sequence across compartments. Our data are consistent with a model in which α5-subunit containing receptors (red) mediate the early phase of postsynaptic memory storage, potentially by leading to elevated calcium flux (not addressed in this study) at individual postsynaptic densities (see discussion and Supplementary Fig. 9 showing separated PSDs and analyses concerning input specificity). Concurrent events see changed dynamics of the α2 receptor (blue). Nicotinic receptor subunits hereby potentially interact with adaptor proteins to bind to Dlg. Importantly, we identify elevated α2 subunit dynamics in the context of associative (M4; KC (green) and dopaminergic neuron (orange) activation needed for memory formation) and non-associative (α’3 MBONs; odor activates both KCs (green) and DANs (purple)) memory expression. Increased α2 subunit dynamics in both cases are triggered by odor application. At the level of M4, suppressed dynamics (concurrent activation of KC (green) and DAN (orange)), would correspond to postsynaptic depression, while at the level of α’3 MBONs increased dynamics could result in postsynaptic depression. Therefore, different learning rules might govern the incorporation, exchange or stabilization of receptors in or out of synapses. Please see discussion for further details.

## Discussion

Synaptic weight changes are widely recognized as substrates for memory storage throughout the animal kingdom. How synapses adapt in order to change their efficacy during learning has been a focus of attention over the last decades. While it is undisputed that both pre- and postsynaptic mechanisms of memory storage exist in vertebrates, invertebrate memory-related synaptic plasticity has been largely localized to the presynaptic compartment, although some debates exist^1^. The core of the debate boils down to a key question: do vertebrates and invertebrates use similar mechanisms to store memories or are there fundamental differences? A first clear difference appears to be the use of different neurotransmitter systems, glutamate and acetylcholine respectively, in the vertebrate and *Drosophila* learning centers^4^.

### Postsynaptic plasticity in appetitive memory storage

Here, we use the genetic tractability of the *Drosophila* system to directly address postsynaptic plasticity during memory storage in invertebrates. Large amounts of evidence from *Drosophila* so far suggests a presynaptic mode of memory storage^24–28,46^. Moreover, it was demonstrated that block of KCs during learning does not interfere with memory performance^28,30,31^, although some studies blocking KC subsets did find impairments^29,47^ in the context of short-term appetitive memory. Lack of necessity to detect neurotransmitter by the postsynapse in the course of memory storage was interpreted as evidence for a presynaptic storage mechanism. If the receptor does not ‘see’ the signal, it is dispensable for ‘interpreting’ it. Here, we revisited such experiments and found, in accordance with previous studies, only mild, if any, requirement for aversive memory storage. We, however, fully abolished appetitive memories (Fig. 1) by blocking KC output during acquisition, providing a model for postsynaptic plasticity (Fig. 9) that is induced and expressed through distinct nAChR subunits (Fig. 2-8).

Our study hints towards different pre- and postsynaptic storage mechanisms underlying aversive and appetitive memories. It also argues against the assumption that appetitive and aversive memories will necessarily use the same molecular machinery to store information. Findings in the past, predominantly based on investigating aversive memories, have been generalized to learning per se. Indeed, postsynaptic contributions have been ruled out for a synaptic junction required for storage of aversive but not appetitive memories, which is fully consistent with our findings^22^. Interestingly, arguing for a division of appetitive and aversive storage sites, subpopulations of KCs have been implicated in aversive and appetitive memory respectively^48^. Lastly, we do not wish to exclude a potential involvement of postsynaptic plasticity in aversive memory formation per se. On the contrary, it is conceivable that aversive memories also could have an appetitive component (release from punishment). However, our experiments suggest that these influences are not as crucial as for appetitive memories, at least in the context of single trial differential associative learning.

### Postsynaptic plasticity at the KC to MBON synapse

Recent anatomical studies^23,44^ have reported both dopaminergic innervation of presynaptic KC compartments as well as somewhat unexpectedly direct synapses between presynaptic dopaminergic terminals and MBONs. We devised an experiment where we substituted KC input to the postsynaptic MBON compartment through artificial acetylcholine injection, while rendering dopaminergic neurons switchable through optogenetics. A protocol that trained and subsequently tested the synaptic junction between KCs and MBONs, demonstrates that plasticity (represented by a change in calcium responses to acetylcholine injection) was inducible by pairing dopaminergic with postsynaptic MBON activation that lasted beyond the training stage and was observable by mere ‘recall-like’ activation of the system (Fig. 3).

Our proof-of-principle experiments uncovered the ability to potentiate after pairing M6 MBON activation and stimulating a broad population of dopaminergic neurons that convey information on sugar, water or the relative valence of aversive stimuli^5^, while we find postsynaptic plasticity to be required for appetitive memory performance (Fig. 3). However, previous studies looking into ‘natural’ appetitive sugar conditioning uncovered a relative depression in M4 (another MBON of the M4/6 cluster) dendrites, when comparing the responses of the paired (CS+) and unpaired odor (CS-) one hour after appetitive conditioning^5,7,13^. Moreover, we here show that *in vivo* appetitive absolute training depresses subsequent responses to the trained odor (Fig. 5). It is important to note that, here (Fig. 3), for our *in vitro* experiments, we perform global activation of the postsynaptic compartment and not the natural typical coverage of 5 % of input synapses per odor^49^ (that allow for differential conditioning). Induced changes are therefore likely not comparable to the natural settings, where sparse sets of KCs and dopaminergic neurons are active within a tight temporal window. Moreover, we here abolish network contributions (by suppressing active signal propagation), to be able to concentrate on synaptic mechanisms during plasticity induction. Thus, our artificial training (Fig. 3) through global dendritic activation likely does not mirror precise physiological conditions allowing for plasticity of a sparse set of synapses to convey odor-specificity to a memory, and should therefore only be viewed as a proof of principle for postsynaptic plasticity induction per se. However, the relatively small amplitude of potentiation observed actually fits previous^7^ *in vivo* observations. Importantly, because we are using TTX and local training of KC to M6 synapses in our experiments, we are furthermore missing additional disinhibition that *in vivo* is mediated via the GABAergic MVP2 MBON^13,19^. Of note, similar protocols^50^ that involved broad activation of KCs (and thus did not circumvent the presynaptic compartment) have demonstrated comparable plasticity induction at this synaptic junction.

Local acetylcholine application to the MB can also activate calcium transients in dopaminergic presynaptic terminals^51^. Therefore, our protocol could in principle include some dopaminergic contributions already at baseline level. However, control experiments using the paired training protocol in the absence of CsChrimson expression in dopaminergic neurons, do not show any signs of plasticity (Fig. 3 and Supplementary Fig. 3). Moreover, it has been previously demonstrated that, to actually release dopamine from the presynaptic terminal, a coincident signal via carbon monoxide is required^52,53^. Therefore, an unwanted activation of dopaminergic neurons in our experiments is unlikely.

It should also be noted that M4, which shows depression (Fig. 5), and M6 have common but also distinct physiological roles, for instance during aversive memory extinction^13^. Besides that, different temporal requirements for M4 and M6 memory expression have been reported^11^. It is therefore possible that physiological changes in the context of appetitive learning led to different plasticity profiles in M4 and M6 neurons respectively, or that initial potentiation over time can be reverted to depression. As noted above, MBON drive is bidirectionally modifiable and has the propensity to both potentiate and depress^7,11,17,26^.

We did not observe plasticity after paired training following α2 knockdown *ex vivo* (Supplementary Fig. 3). In line with that, our *in vivo* experiments directly establish a connection between learning-triggered synaptic plasticity and the α2 subunit (Fig. 5). Appetitive training induced synaptic depression at the level of the α’2-MBON dendrites of control animals. In animals where we concurrently knock-down α2 in M4/6 MBONs, this synaptic depression, however, was not observable.

### Nicotinic receptors could follow temporal sequence

Lasting plasticity traces as observed here (Fig. 3) appear to fit the core criteria for long-term potentiation of vertebrate glutamatergic postsynapses^3,54^. Plasticity can be divided into an ‘induction’^3^ period mediated via NMDA receptors and a subsequent ‘expression’^3^ period that requires altered AMPA receptor dynamics^3^. Our findings here lead to a model, where the nicotinic α5 subunit is required for the induction of appetitive memories at *Drosophila* MBONs (Fig. 2, 9). We propose that α5 nAChR subunits (that can form homomeric channels^38,39^) could take on a similar role to NMDARs. α5 would gate the potentiation or depression of synaptic strength influencing the incorporation or exchange of additional receptor subunits or complexes. In line with this, we show that knock-down of α5 subunits interferes with familiarity learning in the α’3 compartment of the MBs: flies no longer form familiarity memories, they react to a novel odor the same way as to a familiar one, ‘as if they had learned that this new odor was familiar before’ (Fig. 7, 8). Moreover, we do not observe α5 subunit dynamics (Fig. 7), whereas knock-down of α5 leads to decreased levels of α2 subunits (Fig. 4), and α2 dynamics are no longer observable when knocking-down α5 in the MBONs of the α’3 compartment (Fig. 7). Thus, we can draw first analogies to glutamatergic systems governing plasticity in vertebrates. Whether more core criteria are met for the comparison of invertebrate and vertebrate plasticity systems, further depends on whether the here observed receptor dynamics will actually translate to exo-/endocytosis of postsynaptic receptors or lateral diffusion of receptor subunits along the MBON dendrites. Our established system should provide the means to investigate this further in the future.

Interestingly, high levels of dendritic activation after α5 knock-down are translated to reduced axonal calcium transients^4^, effectively leading to decreased signal transduction within the MBON. Of note, MBONs do not appear to exhibit prominent spines on their dendrites^23^ (but see section below: *Are cholinergic and glutamatergic synapses interchangeable?*). Therefore, increased dendritic activation could lead to a change in membrane resistance and result in synaptic interference.

### α2 subunit-positive receptors mediate memory expression

We also find that later forms of appetitive memory expression require both the α2 and α1 receptor subunits (Fig. 2). A recent study^37^ has demonstrated that, when expressed heterologously, these subunits can co-assemble to form heterodimers with β subunits, which, depending on the precise composition of these channels, can harbor different properties, potentially reminiscent to AMPAR^55^. However, MB distribution profiles of α1 and α2 subunits do not match completely, for instance at the level of the γ5 or α’2 compartments (Fig. 4), indicating that they could also partake in different or independent receptor configurations. It would be worthwhile to compare receptor localization with single cell sequencing results^8^.

Importantly, α2 subunit knock-down at M4/6 MBONs does not affect immediate appetitive memories, but later stages of memory (Fig. 2). We show in a complex neuropile, that *in vivo*, on a time scale of 10 to 20 minutes after learning, exposure to odor induces changed α2 receptor subunit dynamics (signal recovery after photobleaching) that are suppressed by simultaneous dopamine exposure (‘learning’, Fig. 6). Therefore, changes in α2 dynamics could match the temporal profiles of memory expression and consolidation processes taking place in an early phase following training (the first 20 minutes) to establish longer-term memories (3 hours). It should be noted that, whether dopamine pairing would also lead to a reduction of α2 levels over time, potentially explaining synaptic depression, cannot be assessed with our experimental settings at this time, but would appear as a valid possibility.

We show that familiarity learning can take place when knocking down α2 nAChR subunits in α’3 MBONs in principle (Fig. 8), however, at clearly decreased efficacy and only after several trials. We speculate that the observation of memories still being expressed *per se* in this context, could be explained by redundancies with α1 or other subunits (but see heterogeneous localization and enrichment in different MB compartments, Fig. 4). Redundancies could also explain why we partially observe functional phenotypes after knock-down of individual subunits, but only moderate structural changes. We also would like to point out that subunits we did not identify as absolutely required for memory expression (Fig. 2) in this study could nonetheless partake in distinct phases of plasticity processes.

Similar to AMPAR, localization of α2, but not α5 subunits, depends on Dlg, the orthologue to vertebrate PSD-95 and PSD-93. Interestingly, the latter has been implicated in structural integrity of cholinergic nicotinic receptor arrangements^59^ in vertebrates.

### α2 dynamics as general plasticity mode?

In the context of both familiarity learning and appetitive conditioning, odor exposure induces increased α2 subunit dynamics (Fig. 6, 7) accompanying postsynaptic depression^7,15^ (Fig. 7), while not or mildly affecting α5 subunits (for familiarity learning). Therefore, the same basic mechanisms, odor-induced α2 receptor dynamics, seem to express two opposed plastic outcomes in the context of associative and non-associative memories and contribute to different learning rules across MB compartments^22,56^. We speculate that α2 dynamics induced by odor in the M4/6 dendrites could be reminiscent of dark currents in the vertebrate visual system^57^ allowing for rapid adaptation with low levels of synaptic noise. Receptor exchange at the level of M4/6 dendrites would actually take place when no associations are formed and stalled when dopaminergic neurons (triggered by sugar) are simultaneously active with KCs (triggered by odor). Indeed, repeated OCT stimulation led to a facilitation of calcium transients (potentially corresponding to an increase of receptor incorporation, Fig. 5-6), while depression (in this case likely to be mediated by removal of receptors, but see above) is triggered by paired training (Fig. 5). In contrast, at the level of the α’3 compartments, odor activates both MBONs and dopaminergic neurons. Here, the plasticity rule would be reversed. Synaptic depression is accompanied by actively changing the receptor composite. We speculate that such plasticity could function reminiscent of mechanisms observed for climbing fiber-induced depression of parallel fiber to Purkinje cell synapses^58^. However, whether increased dynamics can be translated to more incorporation or removal of α2-type receptors, or depending on the plasticity rule both, will require high resolution imaging experiments in the future.

### Are cholinergic and glutamatergic synapses interchangeable?

Our study fuels the question of how unique properties of individual neurotransmitter systems at synapses are. While dopamine signaling is remarkably conserved between invertebrates and vertebrates, cholinergic and glutamatergic systems appear, now more than before (with this study), somewhat interchangeable. While vertebrates (but also evolutionarily distant *C. elegans*) for instance use acetylcholine at the neuromuscular junction and store memories predominantly at glutamatergic synapses, it is the other way around in *Drosophila* and other invertebrates, such as *Sepia*^4–6,60,61^. Now we show that, at cholinergic synapses, α5 and α2 subunits, at least to a certain extent, behave in a potentially comparable way to NMDARs and AMPARs at glutamatergic synapses during postsynaptic plasticity which underlies memory storage. In this context, we offer several lines of evidence that invertebrates utilize postsynaptic plasticity during memory storage and not as previously assumed rely on presynaptic mechanisms only.

We therefore propose that, across phyla, postsynaptic plasticity, with the propensity to store memories and adapt network function plastically, can take place regardless of neurotransmitter identity.

One key difference between the dendritic arbors of the MBONs analyzed in this study compared to dendrites of glutamatergic neurons in vertebrates, is a lack of anatomical spines (Supplementary Fig. 9a). Without spines, how can input-specificity be preserved at MBON postsynaptic densities? KC output to MBON input analysis of the recently published fly hemibrain connectome^62^ (neuprint.org) suggests that, at the ultrastructural level, MBON postsynaptic densities are separated spatially (Supplementary Fig. 9a). Compartmentalization could therefore be mediated by, for instance, biochemical separation of PSDs. Importantly, input-specific plasticity has been shown to be inducible in non-spiny neurons in vertebrates, with diffusion barriers established, e.g., through calcium buffers, between postsynaptic densities^63^.

Interestingly, our MBON input analysis further revealed that postsynaptic plasticity mechanisms could actually add a layer to promote input-specificity. Indeed, we find that single presynaptic KC release sites that innervate MBON dendrites can also target other MBONs and/or other postsynaptic targets simultaneously (Supplementary Fig. 9b-f). Plasticity confined to single postsynaptic densities innervated by a KC terminal could therefore change the weight of transmission for one target (e.g., MBONs involved in memory storage, such as M4^7^), while not changing the weight of the connection to other targets (for example MBONs not involved in a specific action or other targets of non-MBON identity, Supplementary Fig. 9g). It should be noted that this architecture does not exclude presynaptic plasticity mechanisms^70^ (for instance following aversive conditioning). Indeed, we would speculate that synaptic connections can be subdivided into distinct compartments on both the pre- and the postsynaptic side, potentially through transsynaptic molecules^61,64^, allowing for fine-tuned and target-dependent changes of parameters within either side of a synapse.

### A molecular plasticity sequence?

Together, we propose a model (Fig. 9) in which α5-subunit containing receptors could mediate the early phase of postsynaptic memory storage and we speculate this could lead to elevated postsynaptic calcium flux (not addressed in this study). Concurrent events see changed dynamics of the α2 receptor. Nicotinic receptor subunits hereby could interact with adaptor proteins to bind to Dlg, reminiscent to what is known for AMPAR^35^. Importantly, we identify elevated α2 subunit dynamics in the context of associative and non-associative memory expression. Increased α2 subunit dynamics in both cases are triggered by odor application. At the level of M4/6, suppressed dynamics would correspond to synaptic depression, while at the level of α’3 MBONs increased dynamics may result in postsynaptic depression. Therefore, different learning rules could govern the incorporation or exchange or mobilization of receptors in or out of synapses. The precise molecular and biophysical parameters underlying these plasticity rules are currently unknown and will need to be addressed in the future. One option could include potential exchange of α2 subunits for a receptor complex with higher calcium permeability.

Our findings are consistent with the current mushroom body skew model^5^, where the summed MBON output will determine an animal’s choice. However, we add an additional layer, already at the MBON input site. Changes do not happen, as previously believed, at the presynaptic compartment only, but potentially at both synaptic compartments. Thus, the power to store (potentially conflicting) information separately at either the pre- or postsynaptic site, equips the system with additional flexibility. How precisely pre- to postsynaptic and post- to presynaptic signaling is regulated will need to be addressed in the future, but will likely involve transsynaptic signaling routes^61,64^. Importantly, the identified modes of postsynaptic plasticity will open avenues for investigations looking into pre-versus postsynaptic contributions during reversal learning, reconsolidation and extinction learning^13,33^.

## Methods

### Fly genetics

Flies were raised on standard food under standard laboratory conditions unless stated otherwise (25°C, 65 %, 12-hour light-dark cycle)^7,40^. Driver lines used were MB011B (Split-Gal4)^10^, MB112C^10^ (Split-Gal4), MB461B^10^ (Split-Gal4), MB027B^15^ (Split-Gal4), R13F02-Gal4^10^, OK107-Gal4^4^, VT1211-GAL4^7^, and R58E02-LexA^26^. We used the following UAS-nAChR^RNAi^ flies^4,51^: Bloomington stock numbers 28688, 27493, 27671, 31985, 25943, 27251 and 25835. Additionally, we used^41,42^ Dlg^S97^-RNAi as well as UAS-Dlg^GFP^, tubP-GAL80^ts^^48^, UAS-Shi^ts1^^48^, 247-dsRed^7^, LexAop-CsChrimson, UAS-GCamp6f^4,13,26^, and UAS-SynaptoPhluorin^65^. Gal80^ts^ flies were raised at 18-20°C and were placed at 32°C 3-5 days before the experiment. Note that complex genotypes did not always permit usage of MB011B for genetical access to M4/6 neurons throughout the manuscript. In that case, in order to reduce genetic complexity, we used VT1211-Gal4.

### Behavior

#### T-maze memory

3-9-day old mixed-sex populations were trained and tested together as previously described^7^. Odors used were 3-Octanol (OCT, Aldrich) and 4-Methylcyclohexanol (MCH, Aldrich) diluted in mineral oil (approximately 1:100 for aversive, 1:1000 for appetitive memory, absolute concentrations were minimally adjusted to prevent odor bias). For aversive protocols, flies were exposed to the CS+ for 1 minute with 12 1.5 seconds long 120 V electric shocks (interstimulus interval: 3.5 seconds) followed by 45 seconds of air, 1 minute of CS-exposure and another 30 seconds of air. Flies were given 2 minutes to choose between the CS+ and CS- in a T-Maze during retrieval in the dark. For appetitive conditioning flies were starved for 20 - 24 hours before the experiment. Flies were exposed to the CS- for 2 minutes. After 30 seconds, flies were exposed to the CS+ paired with sugar for 2 minutes followed by another 30 seconds of air. Performance indices were calculated as described previously^7^. Time of retrieval is stated in the figures. For Shi^ts^ experiments, flies were kept at 32°C 30 minutes prior to and during training and brought to room temperature directly afterwards. Room temperature was approximately 23°C. For Fig. 2 and Supplementary Fig. 2 behavioral data sets from separate experiments were pooled. Note that ‘screening hit’ data displayed in Fig. 2a,b and Supplementary Fig. 2a,b were replotted to allow for comparison of genotypes with the corresponding genetic controls in Supplementary Fig. 2i-m.

#### Familiarity learning

Familiarity training was essentially performed as described before^15^ with slight adjustments. Flies were covered in yellow dust (Reactive Yellow 86, Fisher Scientific)^15^ and placed in a cylindrical custom designed chamber. To ensure a constant air stream we placed the chamber between an air and a vacuum pump (800ml/min). Air permeable cotton wool was used to close the open ends of the chamber. The air supply was either connected to pure mineral oil or MCH diluted in mineral oil at a concentration of 1:50. For switching between odor and mineral oil, a clamp was manually opened and closed. Video recording was performed at 26 frames per second. For recordings and analyses we used Python (v3.6) in Anaconda Jupyter Notebook environment.

### Imaging

#### Confocal single photon imaging and receptor quantification

##### Fixed explant brain imaging

Brains were dissected on ice, fixed in 4% paraformaldehyde (Sigma) for 20 minutes and placed in PBST (0.1% Triton) for 30 minutes followed by washing with PBS for 20 minutes twice. Vectashield was used as mounting medium. Flies were 2-8 day-old females raised at room temperature.

##### Recording endogenous fluorescence

Imaging was performed using a confocal single photon inverse microscope (Leica SP5/STED) equipped with a 64x oil objective. Laser power and gain were adjusted between experiments, making normalization of the signals necessary. Values for the heatmap in Fig. 4 were normalized to the mean MB fluorescence value to ensure comparability. Voxel size was (height x width x depth) 123 nm x 123 nm x 500 nm. ROIs were drawn manually in ImageJ using the 247- dsRed channel for orientation (Fig. 4a). Heat maps were created in Microsoft Excel. For quantifications following knock-down, the γ5 compartment was normalized to γ4 ((γ5- γ4)/ γ4), and the β’2 to the β’1 compartment ((β’2- β’1)/ β’2) of the same animal. Each ‘n’ corresponds to one hemisphere.

#### In vivo two photon imaging of receptor dynamics

Fluorescence recovery after photobleaching (FRAP) experiments were performed *in vivo*. 2-8 day-old flies were anesthetized on ice and mounted in a custom-made chamber. The head capsule was opened under room temperature sugar-free HL3-like saline, and legs were immobilized with wax^7^. Sugar-free HL3-like saline containing 30 units of Papain (Roche) was applied to the head capsule for 8 minutes to digest the brain’s glial sheath and facilitate removal. Images were acquired using a multi photon microscope (Nikon) with a 25× water-immersion objective, controlled by Nikon NIS Elements software. Diluted odors (MCH or OCT in mineral oil 1:1000) were delivered on a clean air carrier stream using a 6-channel delivery system (CON electronics). The flies were subjected to experimental conditions including either no odor (air), odor only, odor paired with local dopamine (10 mM) injection via a micropipette, or local dopamine injection only (see Fig. 6b and Supplementary Fig. 6b for experimental protocol schematics). Photobleaching was accomplished using focused, high intensity laser exposure for ∼1 minute. Analysis of fluorescence recovery was performed using FIJI. ROIs were manually selected and the percent recovery fluorescence was calculated by subtraction of the post-bleaching baseline fluorescence and division by the pre-bleaching baseline fluorescence. The percent recovery values were log-transformed and a linear mixed effects model without intercept was fit using the interaction between condition and time as fixed effect - to determine condition-specific differences of the recovery kinetics - and time as random effect (R package lme4). A linear mixed effect model was used to appropriately model repeated measures within animals. Significance of recovery of individual conditions was assessed using the regression coefficients of the condition-time interaction of the linear mixed model. Differences of recovery between pairs of conditions were tested using pairwise comparisons of estimated marginal means of the linear mixed model (R package emmeans). Correction for multiple pairwise comparisons was performed using Tukey’s method.

#### *In vivo* confocal single photon imaging of receptor dynamics and calcium transients

3-4 days after eclosure, female flies were prepared as described above and imaged. Imaging was performed using a SP5 single-photon confocal microscope (Leica microsystems). Recording frame rate was 3 Hz. For bleaching high laser power was used focusing on the α’3 compartments for 15-25 seconds. The baseline was recorded after bleaching, immediately before fixed inter-stimulus interval-training^15^. OCT was presented ten times for a second with a six second pause in between. Odor delivery (CON electronics) was controlled by the Leica acquisition software. After training, the same brain plane was recorded for 10 seconds with a pixel size of 200 nm in time intervals of 0, 5, 10, 15, 20, 30, 60 minutes after training. For control experiments air only was delivered to the chamber. Images of the same time interval recordings were averaged and processed in ImageJ. Gaussian blur (σ = 0.5) was applied for smoothing and ROIs were selected manually.

#### *In vivo* two photon calcium imaging

To measure odor responses, female 3-6-day old flies expressing UAS-GCaMP6f and UAS-RNAi to α2 or α5 at the level of M4/6 were tethered under the multiphoton microscope (Femtonics), essentially as described before^7,66^. 5 alternating 1 second OCT and MCH puffs were applied with 30 seconds in between each presentation. Fluorescent signals were recorded from dendrites in the β’2 MB compartment using MESc software (Femtonics) at a frame rate of roughly 31 Hz. ROIs incorporating the dendritic arbors were manually drawn. Data was processed using a Savitzky-Golay filter. Further analysis was performed using Matlab. For absolute training, following protocol was applied: after initial testing for odor responses, flies were exposed to odor puffs (MCH) twice with a 30-second gap between the applications. Corresponding odor responses were averaged. Training consisted of odor application while the fly fed on a sucrose droplet provided by a custom-made feeding arm^67^. After a two-minute break, two odor puffs with a gap of 30 seconds were applied. Again, corresponding odor responses were averaged. Areas under the curve (AUCs) were calculated using Matlab and the first 4 seconds following odor onset were analyzed in order to cover entire responses. The AUCs pre- and post-training were normalized to the mean pre-training values of a group respectively. The averaged test responses pre-training were compared to the average post-training responses using a paired t-test or Wilcoxon matched-pairs signed rank test.

#### Explant brain widefield imaging, neurotransmitter application and optogenetics

##### Postsynaptic plasticity induction

Brains of 3-10 day-old mixed sex flies were dissected on ice. Flies expressed CsChrimson^tdTomato^ under control of R58E02-LexA and UAS-GCaMP6f (and α2 RNAi) under control of MB011B. The head capsule and sheath were removed in carbogenated solution (103 mM NaCl, 3 mM KCl, 5 mM N-Tris, 10 mM trehalose, 10 mM glucose, 7 mM sucrose, 26 mM NaHCO_3_, 1 mM NaH_2_PO4, 1.5 mM CaCl_2_, 4 mM MgCl_2_, 295 mOsm, pH 7.3) with forceps. The brain was subsequently perfused with carbogenated solution containing TTX (2 µM; 20 ml / 10 min flow speed) and imaged using an Olympus MX51WI wide field microscope with a 40x Olympus LUMPLFLN objective and an Andor iXON Ultra camera controlled by Solis software. An Olympus U25ND25 light filter was placed in the beam path to minimize baseline CsChrimson activation. A custom designed glass microcapillary was loaded with uncarbongenated solution containing 0.1 mM acetylcholine and maneuvered to the M6 dendrites. The injection pressure of a P25-1-900 picospritzer was calibrated between 3-8 psi (depending on initial calcium responses). Each local acetylcholine application spanned 15 ms with a 4 s inter-injection interval.

Three pulses of acetylcholine followed by a 2-3 minute break were recorded after which the optogenetic response was assessed by applying 2 seconds red light pulses with an inter-red light-interval of 4 seconds. 3 acetylcholine pulses were recorded followed by either 5 acetylcholine injections, 5 red light pulses or both paired. For paired training both stimuli began simultaneously and the acetylcholine injection lasted for 15 ms (and gave rise to a calcium transients typically lasting > 1 seconds, please see example in Fig. 3), while the paired red light pulse lasts for 2 seconds, allowing for maximal temporal overlap. This process was repeated 5x, with a 4 second break between trials. Following the training trial, 3 final acetylcholine test injections were applied after 1 minute. For analysis, the first of the 3 acetylcholine injections was always discarded because of initial dilution of the capillary tip and the remaining 2 peak intensities were averaged.

All peaks within an experiment were quantified relative to the fluorescence baseline that we calculated for pre and post training acetylcholine responses. Baselines were set independently for each pre- and post-training recording using the polynomial interpolation function in NOSA^68^. For investigating α2 knock-down, only the paired condition was tested.

For controls not expressing CsChrimson, we used VT1211-Gal4 driving UAS-GCaMP6f, instead of MB011B, for technical reasons. This was combined with either expression of R58E02-LexA or LexAop-CsChrimson^tdTomato^. Only paired training was investigated in this context.

##### Excitability of Kenyon cell axons

To test whether KC axons were excited by focal acetylcholine injections at the level of M4/6 dendrite innervation, either UAS-SynaptoPhluorin or UAS-GCaMP6f were expressed under the control of OK107-Gal4. Following the acetylcholine injection experiment, the capillary was exchanged with a capillary containing the same solution with additional 300-400 mM KCl, to evaluate tissue health (not shown for GCaMP6f imaging). To pick up potentially small changes, we increased the injection pressure to 8-14 Psi and the injection time to 150-225 ms (GCaMP6f, 8s inter-injection interval and 3 consecutive injections) and 300-525 ms (SynaptoPhluorin, with an 8s inter-injection interval and 3 consecutive injections).

Images were analyzed using NOSA^40,68^ and GraphPad Prism.

### Statistics

Statistical analyses were performed as stated in the previous methods sections and figure legends. Data were always tested for normality using a Shapiro-Wilk test. If normally distributed, data were analyzed using ANOVA followed by post-hoc test or a (paired) t-test. If not normal, we used a Kruskal Wallis followed by post-hoc test, or a Wilcoxon matched-pairs signed rank test.

### Tagged receptor subunits

All subunits were tagged using CRISPR technology and motifs previously described^69^. Further details will be published separately and can be requested from the corresponding author.

### Connectome analysis

In Supplementary Fig. 9, we analyzed the partial connectome of the female adult fly brain (hemibrain v1.2.1)^62^ using the neuprint-python package (https://github.com/connectome-neuprint/neuprintpython). To investigate the synaptic relationship between KC presynapses and MBON postsynaptic sites, we first identified KCs with the status ‘Traced’ in the connectome. Second, for each MBON of interest, we identified the relevant KCs connected to it (which is a subset of the count in the previous step). Third, for each KC identified, we selected each presynaptic terminal (x,y,z locations of synapses) connected to the MBON of interest, and for each of these presynapses, we identified all synaptic partners residing on the postsynaptic side. Fourth, for the MBON of interest, we counted the number of postsynaptic connections per individual presynapses (that also contain the specific MBON, Supplementary Fig. 9 d,e,f). Finally, we identified the composition of all neurons identified as co-postsynaptic partners of KC to MBON synapses.

## Data availability

The datasets generated during and/or analysed during the current study are available from the corresponding author on reasonable request.

## Code availability

The code used in this study is available at https://github.com/jagannathancolabs/2022_postsynapticplasticity

## Acknowledgements

We thank Anatoli Ender, Johannes Felsenberg, Davide Raccuglia, Lisa Scheunemann, Stephan Sigrist, Uli Thomas and Scott Waddell for comments on the manuscript, Stephan Sigrist and Uli Thomas for reagents, the Janelia and Vienna fly projects, and the Bloomington Stock Center and VDRC for fly lines as well as Daisuke Hattori and Yoshi Aso for help with the familiarity experiments. Multiphoton and single photon confocal imaging was partially performed using microscopes of the AMBIO and NWFZ core facilities of the Charité.

## Author contribution

Conceptualization, C.P., M-M.H., E.R., S.R., D.L., Y-C.C., D.O., Investigation, C.P., M-M.H., E.R. D.L., R.S.-G., C.R., S.R., Y-C.C., I.B., T.F., G.L., S.R.J.; Resources, D.O.; Writing, D.O., C.P.; Instrumentation: JR; Comments, M-M.H., E.R. D.L., R.S.-G., S.R., Y-C.C., S.R.J..

## Supplemental Information

**Supplementary Figure 1:**
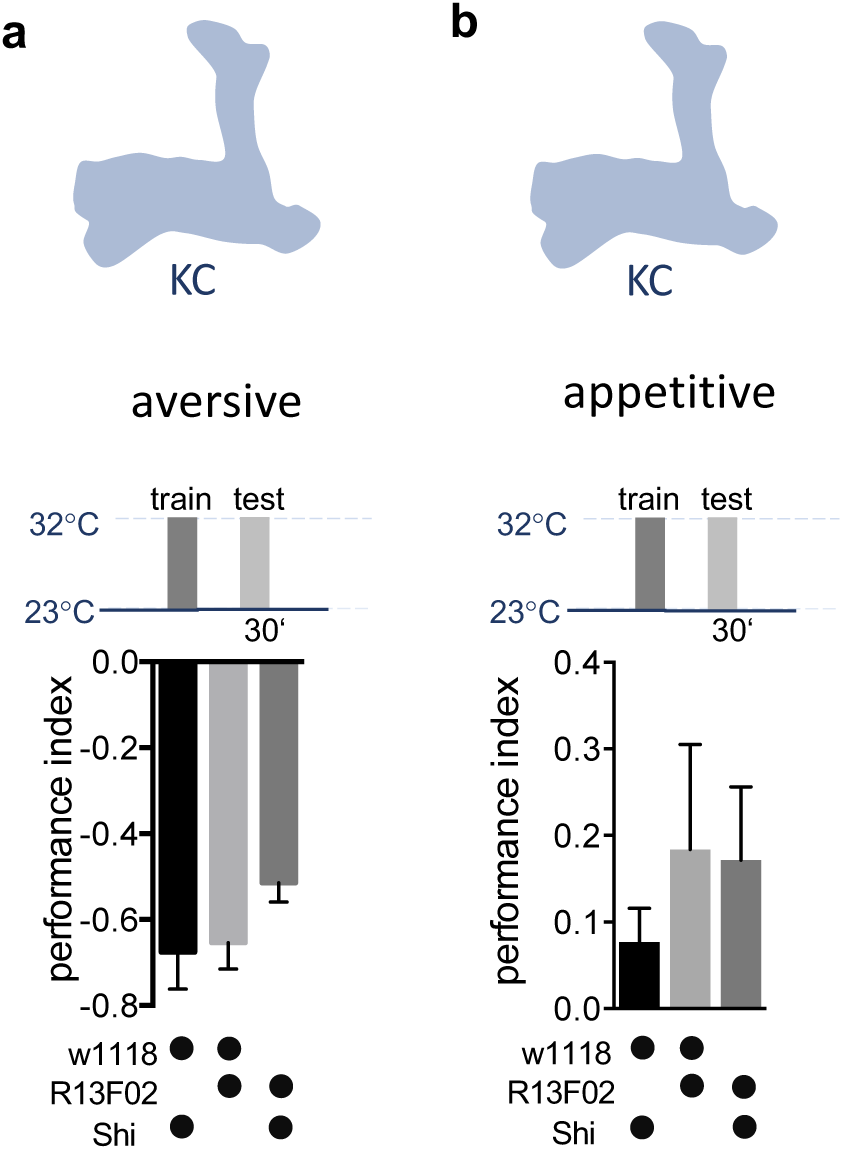
Permissive temperature controls accompanying Fig. 1. **a)** Permissive temperature control for experiments shown in Fig. 1a. 30 min aversive memory performance when training at 23°C (driver line R13F02-Gal4). Bar graphs: mean ± SEM; n = 7 – 9; Kruskal-Wallis followed by Dunn’s test (p = 0.08). **b)** Permissive temperature control for experiments shown in Fig. 1b. 30 min appetitive memory performance when training at 23°C (driver line R13F02-Gal4). Bar graphs: mean ± SEM; n = 6 – 8; Kruskal-Wallis followed by Dunn’s test (p = 0.7).

**Supplementary Figure 2:**
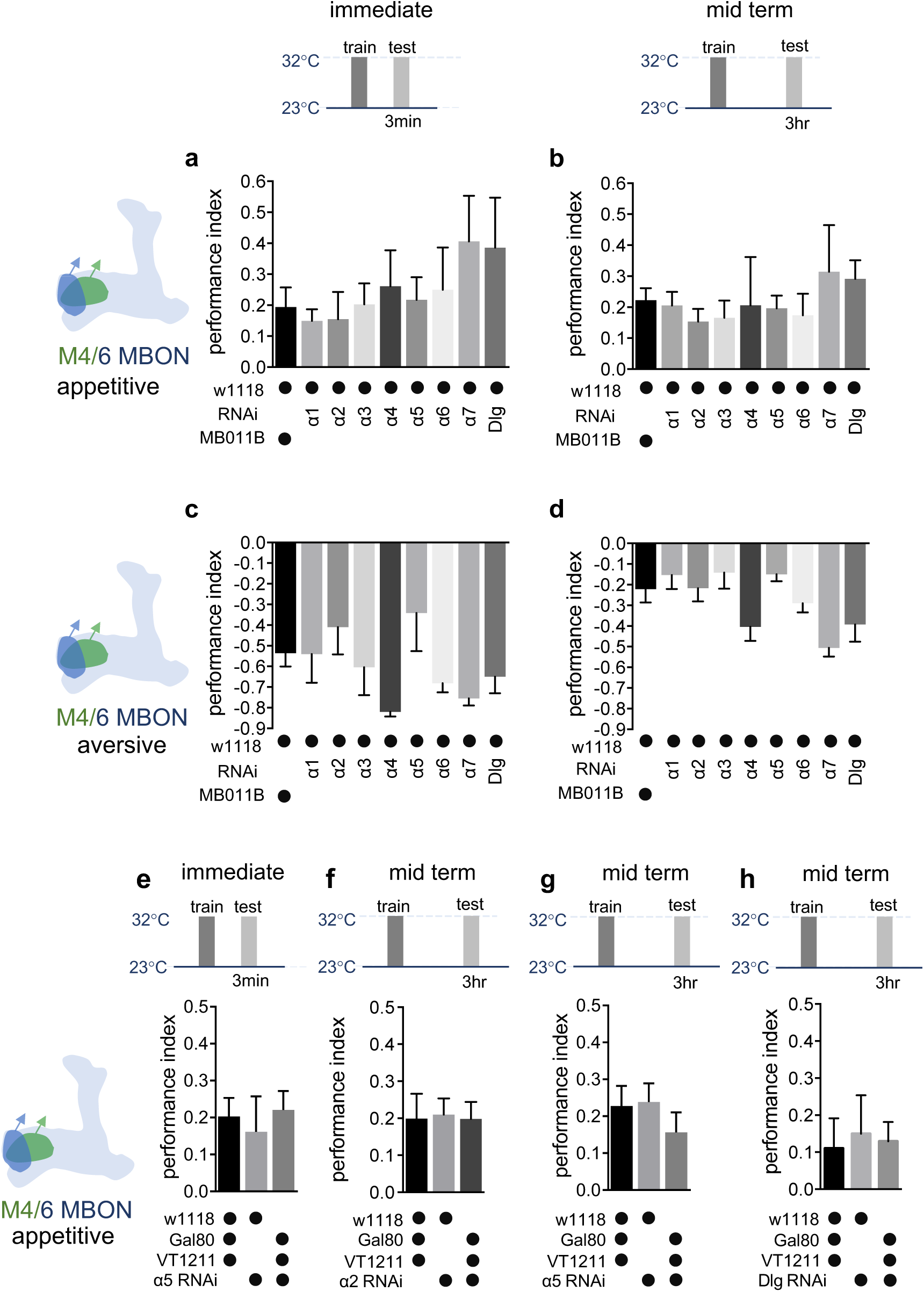
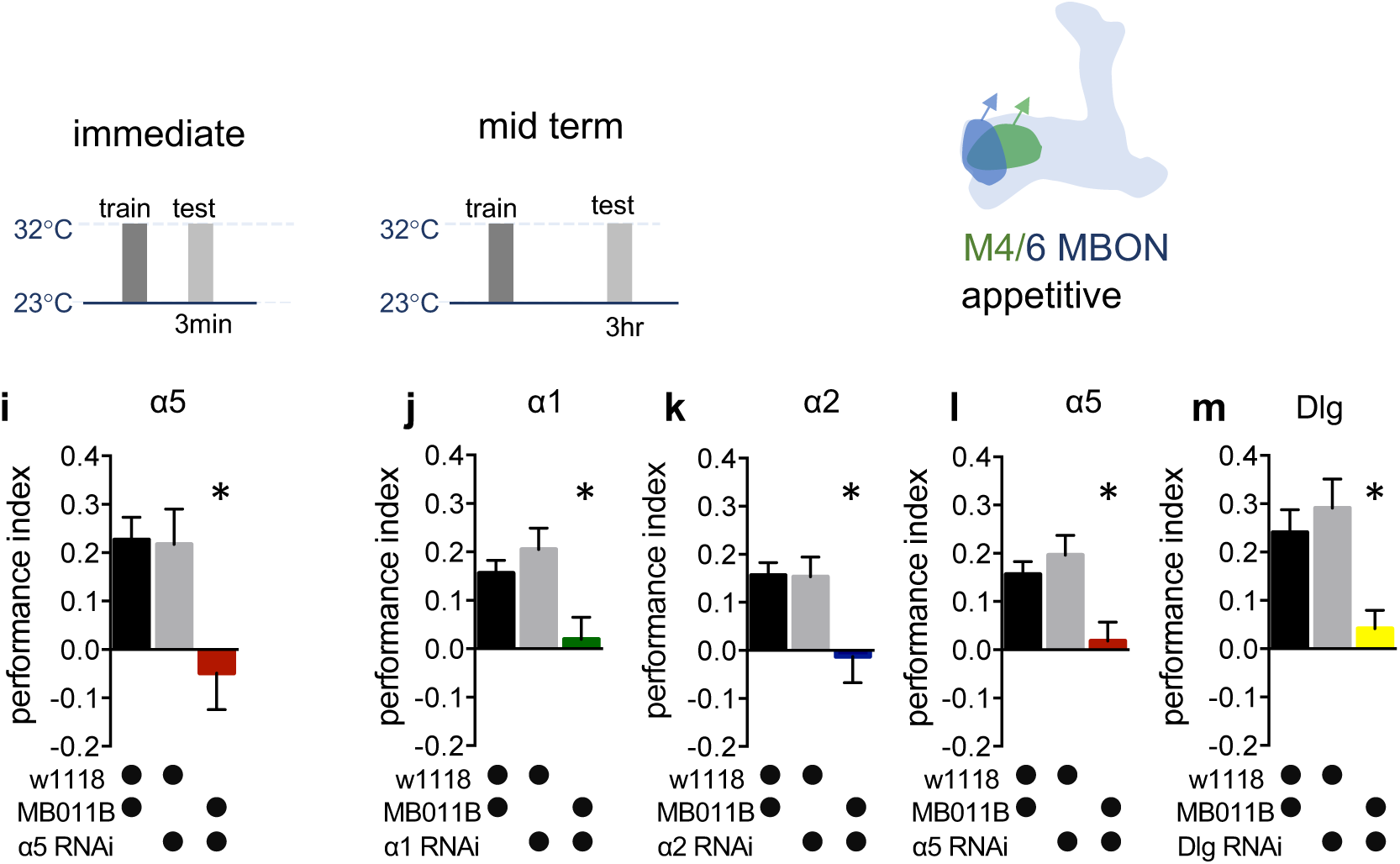
Genetic controls and alternate data display of data presented in Fig. 2. **a)** Immediate appetitive memory is not impaired in genetic control groups. Bar graphs: mean ± SEM; n = 7 – 11, for controls: n = 16, one-way ANOVA followed by Tukey’s test (p > 0.05). **b)** 3-hour appetitive memory is not impaired in genetic control groups. Bar graphs: mean ± SEM; n = 7 – 33, for controls: n = 41, one-way ANOVA followed by Tukey’s test (p > 0.05). **c)** Immediate aversive learning is not impaired in genetic control groups. Bar graphs: mean ± SEM; n = 6 – 10, for controls: n = 13, Kruskal-Wallis followed by Dunn’s test (p > 0.05). **d)** 3-hour aversive memory is not impaired in genetic control groups. Bar graphs: mean ± SEM; n = 6 – 10, for controls: n = 15, Kruskal-Wallis followed by Dunn’s test (p > 0.05). **e)** RNAi knock-down of the α5 subunit in M4/6 MBONs (driver line VT1211-Gal4) is suppressed using Gal80^ts^. Immediate memories is not impaired. Bar graphs: mean ± SEM; n = 9; one-way ANOVA followed by Tukey’s test. **f)** RNAi knock-down of the α2 subunit in M4/6 MBONs (driver line VT1211-Gal4) is suppressed using Gal80ts. Mid term memory is not impaired. Bar graphs: mean ± SEM; n = 9 – 11, one-way ANOVA followed by Tukey’s test. **g)** RNAi knock-down of the α5 subunit in M4/6 MBONs (driver line VT1211-Gal4) is suppressed using Gal80ts. Mid term memory is not impaired. Bar graphs: mean ± SEM; n = 10 – 11, one-way ANOVA followed by Tukey’s test. **h)** RNAi knock-down of Dlg in M4/6 MBONs (driver line VT1211-Gal4) is suppressed using Gal80ts. Mid term memory is not impaired. Bar graphs: mean ± SEM; n = 8 –10, one-way ANOVA followed by Tukey’s test. (**i-m**) Alternate display (see Methods for details) for appetitive memory experiments shown in Fig. 2a,b: **i)** Immediate appetitive memories are impaired following RNAi knock-down of the α5 nAChR subunit in M4/6 MBONs (driver line MB011B). Bar graphs: mean ± SEM; n = 9 – 21, one-way ANOVA followed by Tukey’s test (p < 0.05), * = p < 0.05. **j)** 3-hour appetitive learning is impaired by RNAi knock-down of the α1 nAChR subunit in M4/6 MBONs (driver line MB011B). Bar graphs: mean ± SEM; n = 25 – 51, Kruskal-Wallis followed by Dunn’s test (p < 0.05), * = p < 0.05. **k)** RNAi knock-down of the α2 nAChR subunit in M4/6 MBONs (driver line MB011B) impairs 3-hour appetitive memories. Bar graphs: mean ± SEM; n = 27 – 51, Kruskal-Wallis followed by Dunn’s test (p < 0.05), * = p < 0.05. **l)** 3-hour appetitive learning is impaired by RNAi knock-down of the α5 nAChR subunit in M4/6 MBONs (driver line MB011B). Bar graphs: mean ± SEM; n = 27 – 51, one-way ANOVA followed by Tukey’s test (p < 0.05), * = p < 0.05. **m)** RNAi knock-down of Dlg in M4/6 MBONs (driver line MB011B) impairs 3-hour appetitive memories. Bar graphs: mean ± SEM, n = 13 – 31; one-way ANOVA followed by Tukey’s test (p < 0.05), * = p < 0.05. Note that data in panels a-b and i-m were replotted to allow comparison between all genotypes.

**Supplementary Figure 3:**
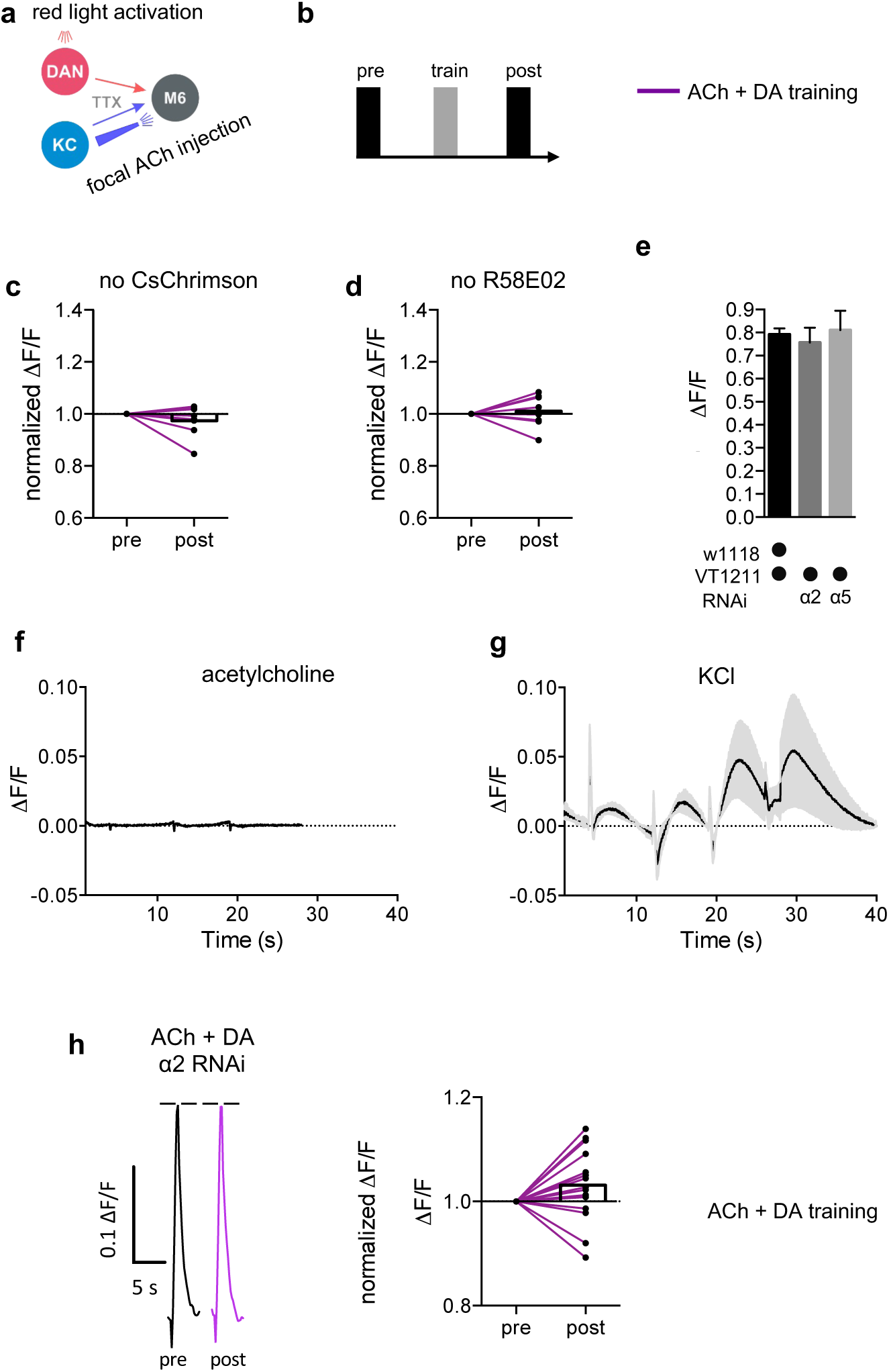
Control experiments and non-normalized data display for Fig. 3. **a)** Connectivity scheme of MB output synapses. Also shown in Fig. 3. **b)** Training scheme. **c)** Control experiments for Fig. 3 f-h. Paired training protocol in the absence of LexAoP-CsChrimson. Before-after plots and bar graphs (mean); n = 9, ratio paired t-test. **d)** Control experiments for Fig. 3 f-h. Paired training protocol in the absence of R58E02-LexA. Before-after plots and bar graphs (mean); n = 8, Wilcoxon matched-pairs signed rank test. **e)** GCaMP6f responses following focal ACh injections to the γ5 compartment of the MB are not significantly altered following α2 or α5 knock-down in the M4/6 MBONs (driver line VT1211-Gal4). The bath contains TTX to suppress spontaneous neural activity. Bar graphs: mean ± SEM; n = 12 – 23, one-way ANOVA followed by Dunnett’s test (p > 0.05). **f)** Averaged SynaptopHluorin responses following focal ACh injections (0.1 mM) to the γ5 compartment of the MB using the OK107-Gal4 driver (KCs), n = 8. Shaded area depicts SEM. **g)** Averaged SynaptpHluorin responses following focal KCl injections (300-400 mM) to the γ5 compartment of the MB using the OK107-Gal4 driver (KCs), n = 4. Shaded area depicts SEM. **h)** Left: sample calcium traces in response to ACh injections recorded from M6 dendrites pre (black traces) and post (colored traces) training. Right: peak quantification. Following RNAi knock-down of the α2 subunit in M4/6 MBONs no significant potentiation was detected after paired acetylcholine application and dopaminergic neuron activation. Before-after plots and bar graphs (mean); n = 15, ratio paired t-test.

**Supplementary Figure 4:**
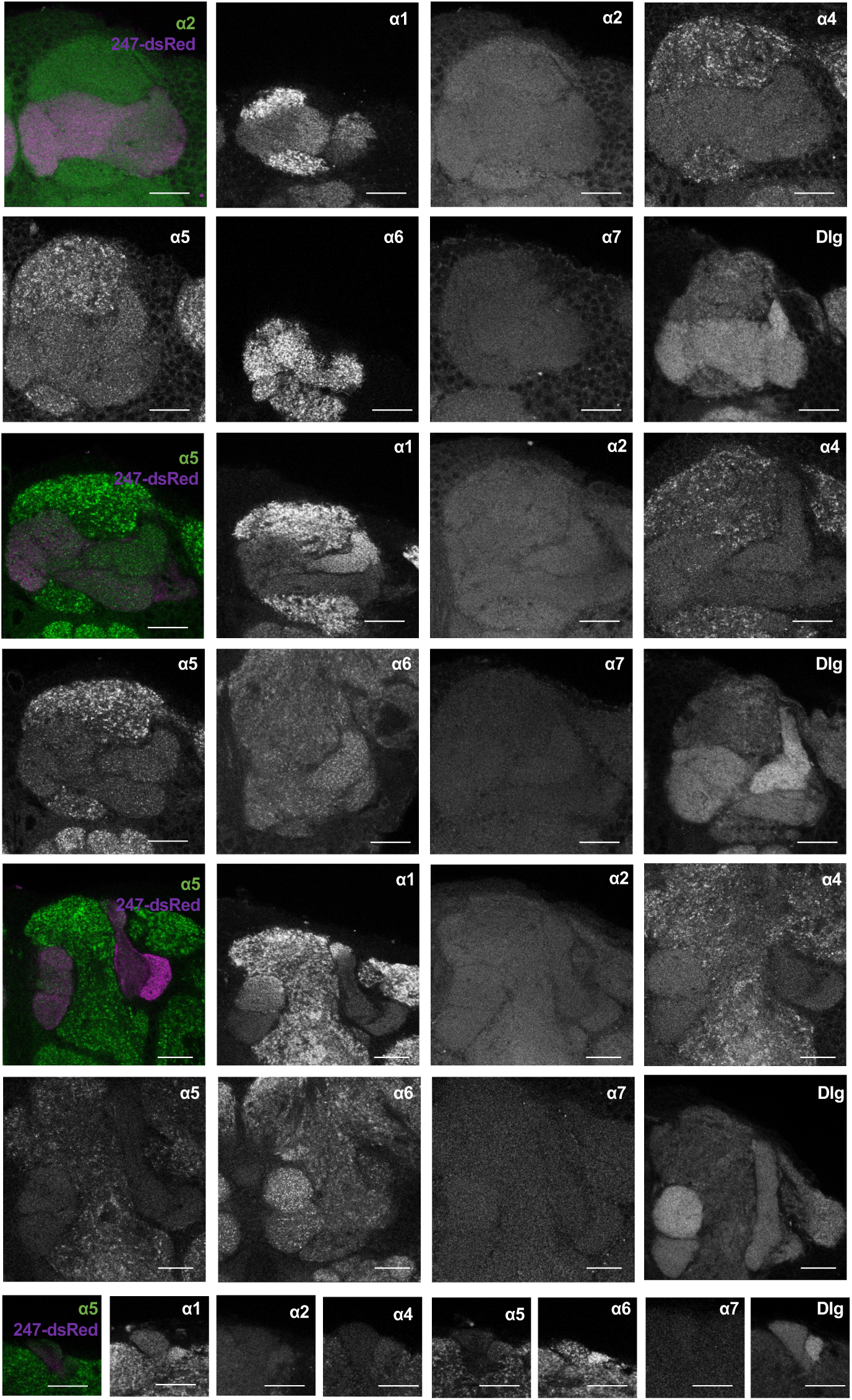
Detailed distribution of α subunits in the MB accompanying Fig. 4. Example image planes of GFP expression at the level of the MB compartments for all α nAChR subunits (except for α3) and Dlg. Pictures are taken from different animals. Scale-Bar: 20 μm. Left: merged image of α subunit signal (green) with MB compartments marked by 247-dsRed (magenta). Top two rows: γ compartments; middle two rows: γ, β’ and β compartments, bottom three rows: α’/β’ and α/β, with bottom row: α’ and α compartments.

**Supplementary Figure 5:**
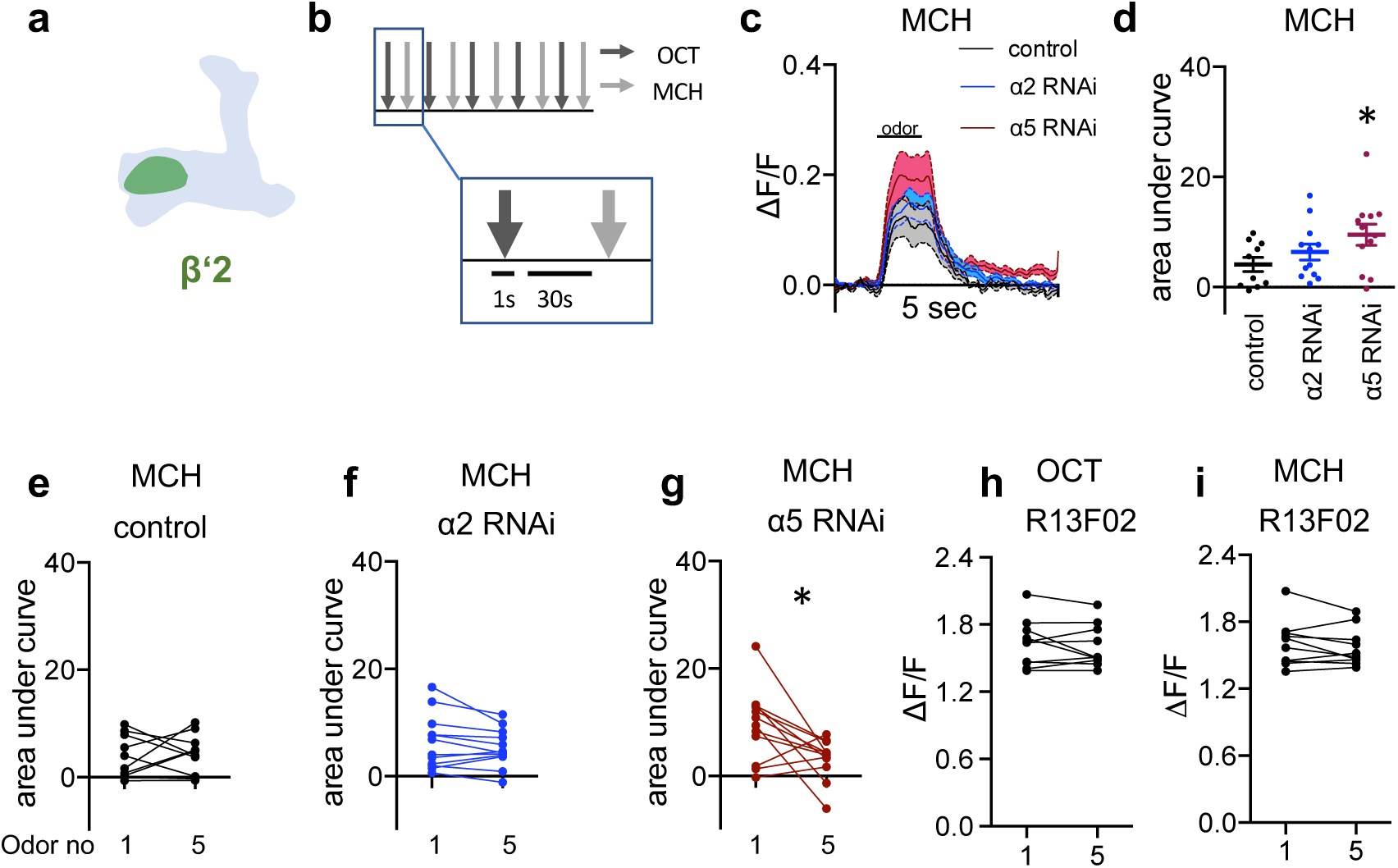
Additional data and display of non-normalized data accompanying Fig. 5. **a)** Scheme indicating the imaging area at the level of the β’2 compartment. **b)** Odor exposure protocol. 5 OCT stimuli were alternatingly administered with 5 MCH stimuli. 1 second odor puffs were separated by 30 seconds breaks. Odor responses are measured using GCaMP6f as in Fig. 5. **c)** Averaged traces of GCaMP6f (calcium) responses to MCH from control (black), α2 (blue) and α5 RNAi (red; driven in M4/6 respectively, driver line VT1211-Gal4) flies. Solid traces are mean, shaded areas SEM; n = 10 - 12. Odor exposure is indicated by bar. **d)** Area under curve quantification of averaged odor responses show significantly elevated odor responses to MCH following α5 knock-down in M4/6 neurons (driver line VT1211-Gal4). Mean ± SEM; one-way ANOVA followed by Dunnett’s test (p < 0.08), * = p < 0.05; n = 10 – 12. **e)** Control flies show no significant increase between the first and the fifth response to MCH. n = 10; paired t-test. **f)** α2 RNAi flies show no difference between the first and fifth odor response to MCH. nAChR subunit RNAi is driven in M4/6 neurons (driver line VT1211-Gal4). n = 12; paired t-test. **g)** α5 RNAi flies show a significant decrease in calcium transients over the course of consecutive odor exposures. nAChR subunit RNAi is driven in M4/6 neurons (driver line VT1211-Gal4). n = 12; Wilcoxon matched-pairs signed rank test; * = p < 0.05. **h)** KCs (driver line R13F02-Gal4) show no significant change in calcium transients over the consecutive OCT odor exposures. Before-after plots. n = 10, paired t-test. **i)** KCs (driver line R13F02-Gal4) show no significant change in calcium transients over the consecutive MCH odor exposures. Before-after plots. n = 10, paired t-test.

**Supplementary Figure 6:**
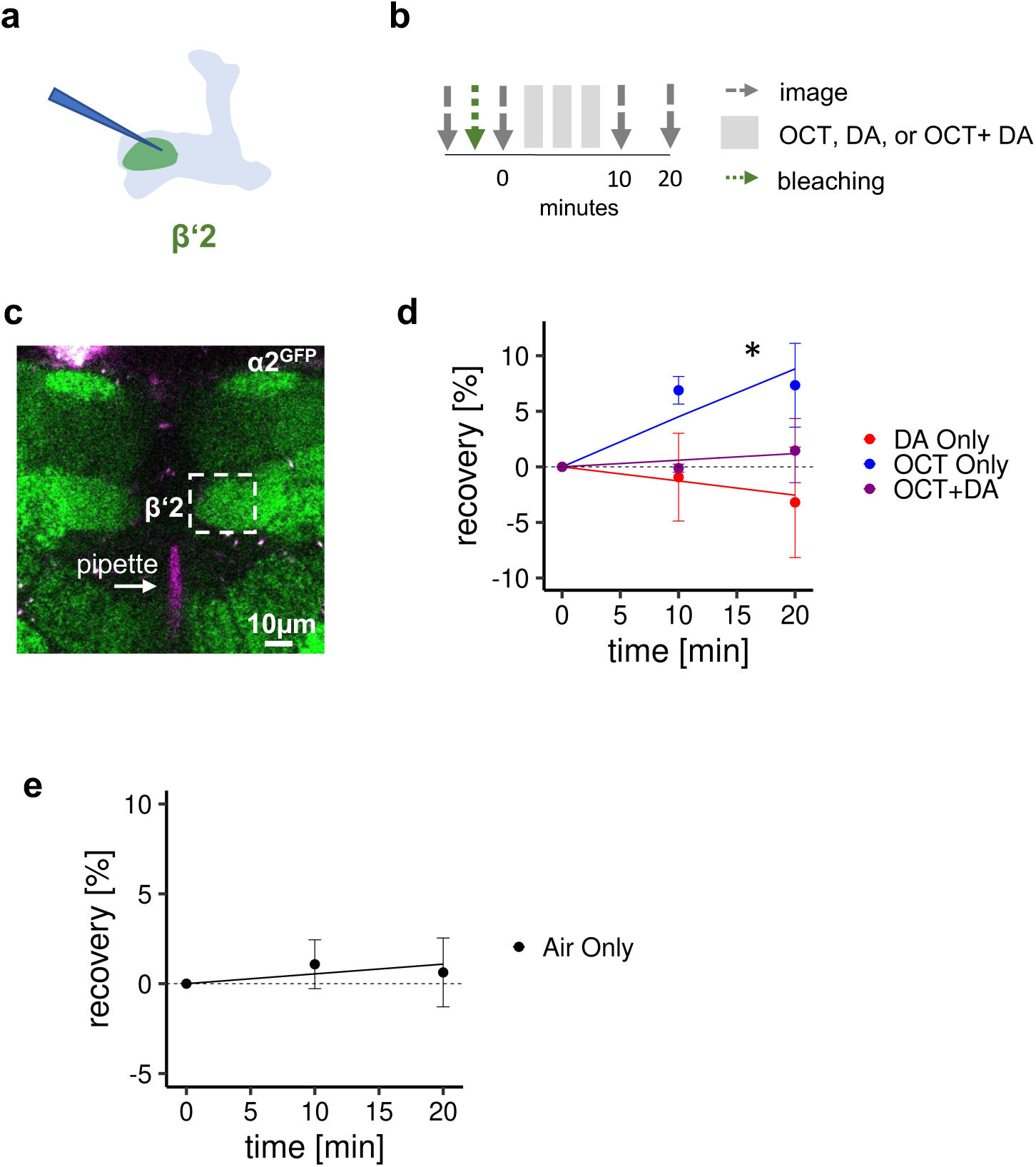
Receptor subunit recovery, accompanying Fig. 6. **a)** Scheme of site of dopamine injection during fluorescence recovery after photobleaching (FRAP) experiments at the level of the KC-MBON synapses of the β’2 compartment. **b)** FRAP experimental protocol. After bleaching, a baseline picture was taken followed by odor presentation or odor presentation simultaneously with dopamine injection or dopamine injection by itself. Fluorescence recovery was monitored at the 10 and 20 minute time points. **c)** Example image of α2^GFP^ expression (the same image is shown here as in Fig. 6); white dashed box shows the β’2 output zone; dopamine injection pipette (with Texas Red) is labelled in magenta. Scale bar: 10 µm. **d)** Regression of fluorescence recovery after photobleaching following OCT exposure (blue line), or OCT exposure simultaneously with dopamine (DA) injection (purple line), or dopamine injection alone (red line). n = 3 –5, t-test, Satterthwaite’s method for approximating the degrees of freedom. *: regression coefficient p < 0.05 in the linear mixed effect model. Only ‘OCT only’ exposure shows recovery that significantly differs from 0. **e)** Control settings for Fig. 6 and the experiments shown in this figure. After bleaching, α2^GFP^ flies were exposed to air only. n = 9, t-test, Satterthwaite’s method for approximating the degrees of freedom. The regression coefficient is not significantly different from 0.

**Supplementary Figure 7:**
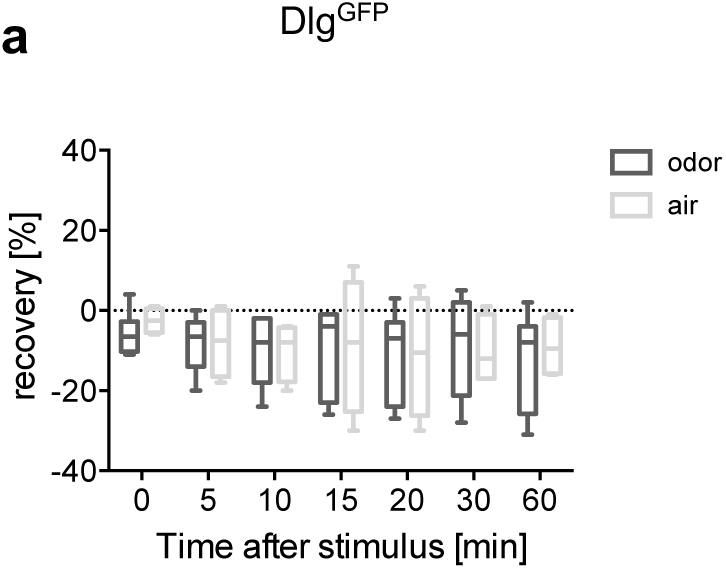
Dlg^GFP^ FRAP, accompanying Fig. 7. **a)** FRAP of Dlg^GFP^ in α’3 MBONs. Dlg^GFP^ did not show significant recovery. Recovery rate is normalized to the baseline recorded after selective bleaching of the α’3 MB compartment. n = 5 - 7; multiple t-tests.

**Supplementary Figure 8:**
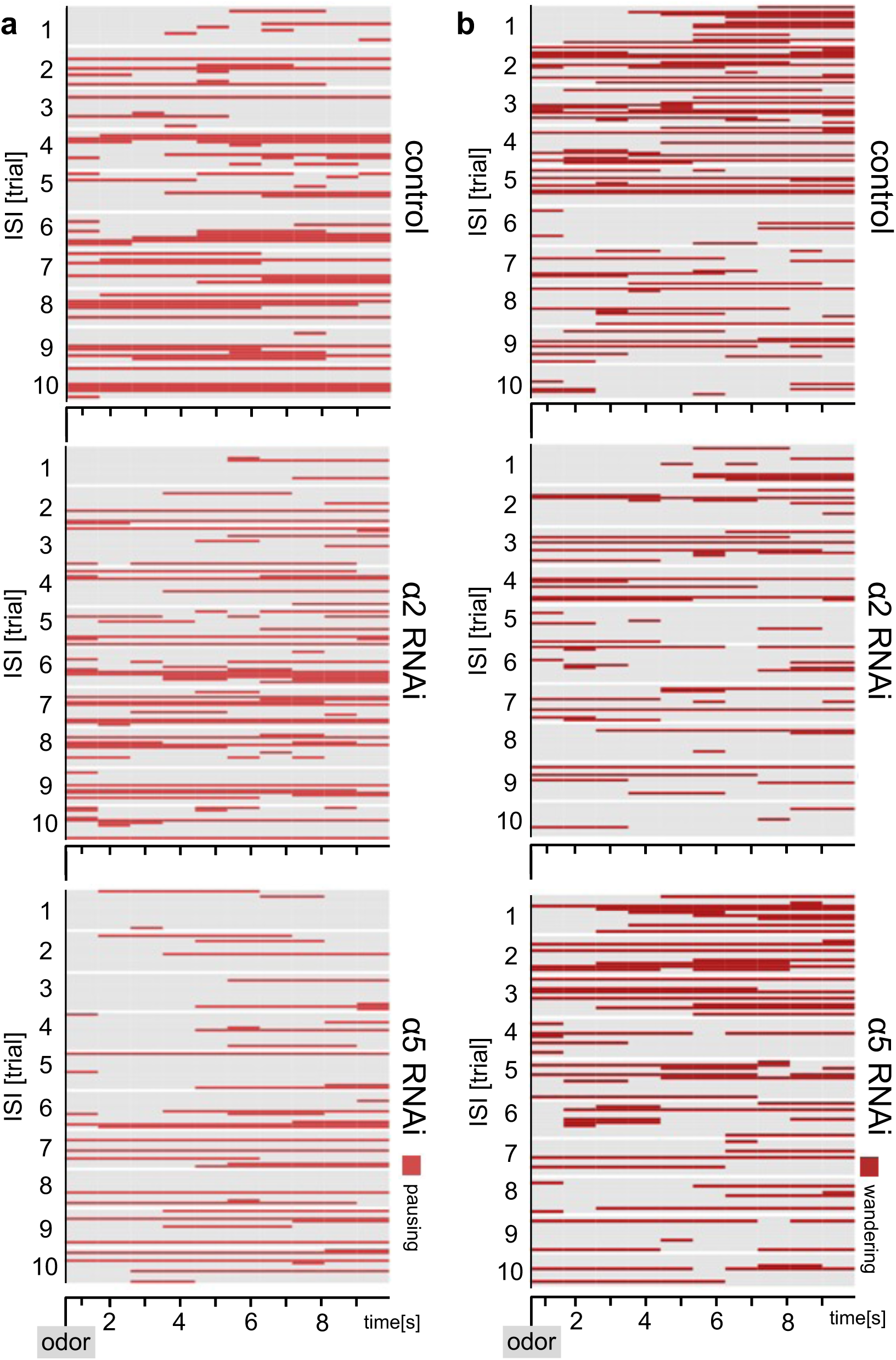
Additional ethograms, accompanying Fig. 8. **a, b)** Ethograms of the behavioral responses of flies shown in Fig. 8 with additional behavioral categories of pausing and wandering (when not grooming). Ethograms show pausing (red, **a**) which is defined as not moving and not grooming or wandering (dark red, **b**) which is defined as moving around in the chamber.

**Supplementary Figure 9:**
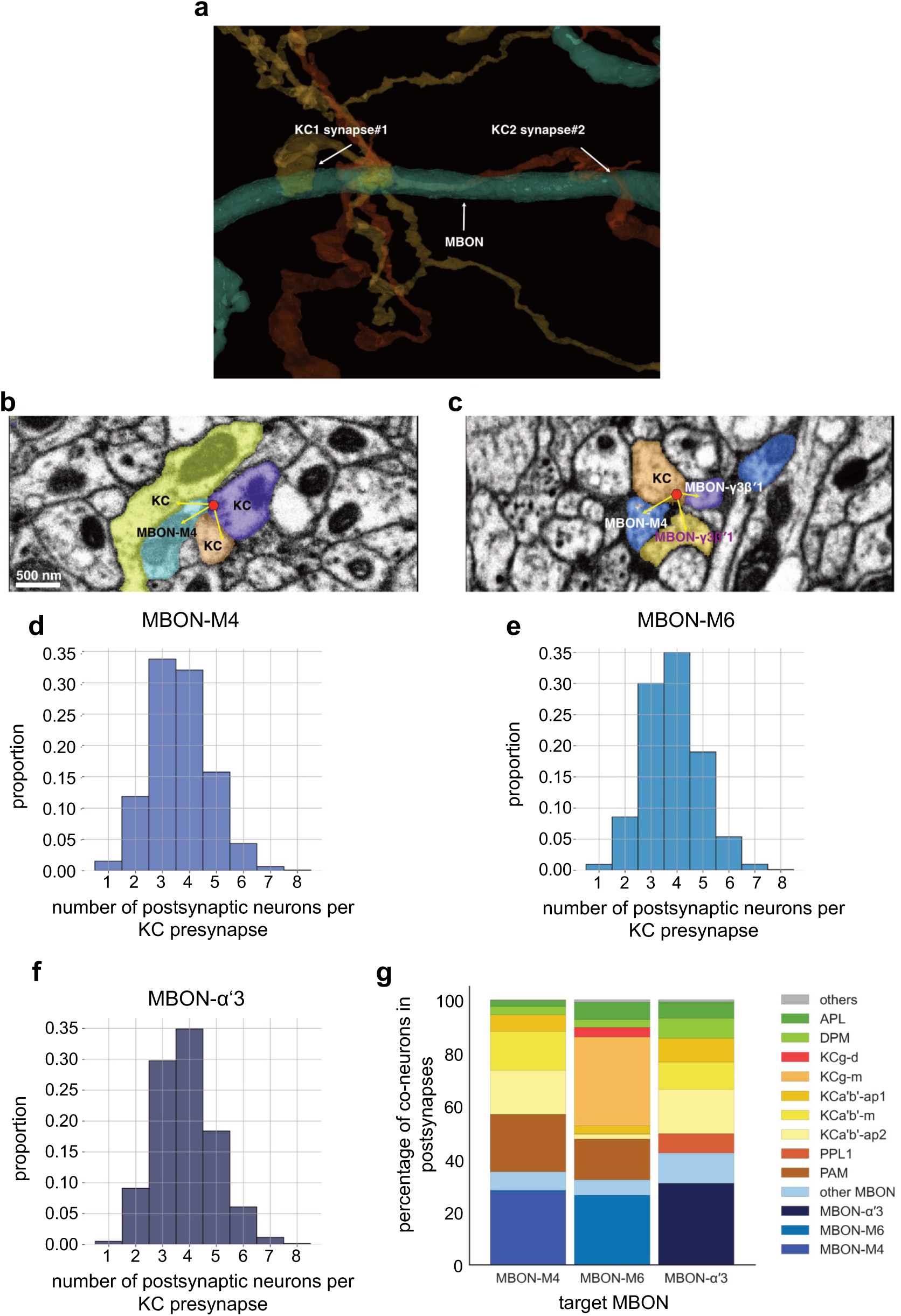
Input analysis of KC to MBON synapses, accompanying Fig. 9. **a)** Reconstructed example synapses from electron microscopic (EM) volume (neuprint.org)^62^: two different KCs connect to the same MBON on the postsynaptic side. **b)** EM image (neuprint.org) showing a KC presynapse simultaneously innervating two other KCs and the M4 MBON. Scale bar: 500 nm. **c)** EM image (neuprint.org) showing a KC presynapse simultaneously innervating the M4 MBON and two sites of another MBON. **d)** Analysis of number of postsynaptic partners for each KC presynapse identified providing input to M4. The histogram shows the distribution of KC synapses to M4 relative to how many postsynaptic partners the KC presynapses have. **e)** Analysis of number of postsynaptic partners for each KC presynapse identified providing input to M6. The histogram shows the distribution of KC synapses to M6 relative to how many postsynaptic partners the KC presynapses have. **f)** Analysis of number of postsynaptic partners for each KC presynapse identified providing input to α’3 MBONs (pooled). The histogram shows the distribution of KC synapses to α’3 MBONs relative to how many postsynaptic partners the KC presynapses have. **g)** Percentage of types of neurons that share a KC presynapse with a given MBON.

